# Amino acid variants at the P94 position in *Staphylococcus aureus* class A sortase modulate substrate binding and enzyme activity

**DOI:** 10.64898/2026.01.18.700168

**Authors:** Noah Cox-Tigre, Mia E. Stewart, Jackson Tucker, Erich G. Walkenhauer, Cooper S. Wilce, John M. Antos, Jeanine F. Amacher

**Author notes:** Corresponding Authors: Jeanine Amacher, Department of Chemistry, Western Washington University, 516 High St – MS9150, Bellingham, WA, 98225, **Email:**, John Antos, Department of Chemistry, Western Washington University, 516 High St – MS9150, Bellingham, WA, 98225, **Email:**.

## Abstract

The surface of gram-positive bacteria is a highly regulated environment with specific attachment of proteins required for viability. Sortase enzymes are cysteine transpeptidases that recognize and ligate substrates to the peptidoglycan layer in these microorganisms, which can be highly pathogenic (e.g., *Staphylococcus aureus, Streptococcus pyogenes,* etc.). As such, sortases represent a potentially novel target for antibiotic development. In addition, the catalytic activity of sortase enzymes is utilized in sortase-mediated ligation (SML) engineering approaches for a variety of uses. In SML experiments, engineered variants of *Staphylococcus aureus* sortase A (saSrtA) are the most widely used enzymes. One of the mutated amino acids in the previously engineered pentamutant (or saSrtA5M) enzyme is P94. Structural analyses of experimental saSrtA structures revealed that P94 interacts directly with Y187 when saSrtA is in its inactive conformation. While saSrtA5M, developed via directed evolution, contains a P94R mutation, we wanted to interrogate this position further and ask if other single P94 mutations may reveal a greater effect on activity and/or substrate specificity. We created 18 P94X mutations (excluding P94C), and tested relative activity using a fluorescence resonance energy transfer (FRET) assay for 4 substrate sequences: LP**A**TG, LP**E**TG, LP**K**TG, and LP**S**TG. We identified several P94 variants that outperformed the single mutant P94R for all peptides tested, including P94A, P94D, P94E, P94G, P94H, P94N, P94Q, P94S, and P94T. We further observed that the reactivity of substrates with variations in the central position of the pentapeptide recognition motif (LP**X**TG) can be sensitive to the identity of the P94X residue. We tested P94A and P94D saSrtA5M variants and found that, depending on LP**X**TG sequence, these variants could outperform saSrtA5M in activity > 3-fold. Finally, we compared saSrtA5M and P94D saSrtA5M in a model sortase-mediated ligation reaction using a LP**K**TG substrate and saw ∼2-fold greater product formation. Taken together, we characterized an important position that modulates substrate access and activity in saSrtA. Furthermore, we argue that future studies which combine rational design and high throughput approaches, e.g., directed evolution, may result in sortase variants with increased SML potential.

## Introduction

The thick peptidoglycan layer that defines gram-positive bacteria, which include pathogens such as *Staphylococcus aureus, Streptococcus pyogenes,* and *Enterococcus faecalis,* provides a protective layer for these organisms.^1^ This surface is a dynamic environment, and acts as a scaffold for proteins that are critical for viability, defense, pathogenesis, and cellular function.^1^ Sortase enzymes, specifically class A sortases (SrtAs), are cysteine transpeptidases localized to the cleavage furrow of dividing gram-positive bacteria, and which covalently attach substrate proteins to the developing peptidoglycan layer.^1,2^ SrtA enzymes were first described in 1999.^3–5^ Even in those first studies, SrtA enzymes were identified as a potential novel target for antibiotic development, work that remains ongoing.^5–19^ In addition to interest in sortase enzymes as antimicrobial targets, sortase-mediated ligation (SML) strategies quickly emerged as a powerful protein engineering approach for *in vitro* and *in vivo* applications, including in nanobody/antibody drug conjugation, developing novel insulin derivatives, as both diagnostic and therapeutic tools for neurodegenerative disease, and in creating multivalent vaccines for SARS-CoV-2, amongst many other uses.^20–29^

The sortase catalytic mechanism is well understood. A core triad of His-Cys-Arg residues (for saSrtA, H120-C184-R197) facilitate the processing of the cell wall sorting signal (CWSS) sequence, LPXTG, where X=any amino acid (L=P4, P=P3, X=P2, T=P1, G=P1’), on a target protein.^30,31^ Recent work from ourselves and others argues that while R197 interacts with and stabilizes the bound LPXTG ligand, it is not directly responsible for catalysis.^26,32–34^ Instead, upon nucleophilic attack of the side-chain thiol of C184 on the carbonyl carbon of P1 Thr, the resulting oxyanion tetrahedral intermediated is stabilized by a conserved Thr immediately preceding the catalytic Cys (in saSrtA, T183) and the backbone amide of the amino acid immediately following the catalytic His (in saSrtA, T121).^26,33^ Overall sortase catalysis follows a ping-pong mechanism, first involving the formation of a thioester-linked acyl enzyme intermediate. Transpeptidation is then completed when a second nucleophilic substrate, typically an N-terminal Gly, attacks the acyl-enzyme intermediate, resulting in covalent attachment of the two substrates via a new peptide bond.^26,31^

Despite the identification of over 10,000 enzymes in the sortase superfamily, SML experiments most commonly utilize engineered variants of the first SrtA discovered, *Staphylococcus aureus* SrtA (saSrtA).^35,36^ Biochemical and structural characterization of a number of SrtA enzymes previously revealed both shared and unique attributes of saSrtA and SrtA enzymes from other organisms. For example, the catalytic domains of SrtA enzymes studied to date share relatively low catalytic efficiencies as compared to other enzymes.^31^ Uniquely, saSrtA and other SrtA enzymes in the *Staphylococcus* genera contain a calcium-binding site, an allosteric activator which initiates a conformational change and is required for full activation of the enzyme.^30,31,36^ Of several SrtA enzymes studied, saSrtA is the most specific for the canonical pentapeptide recognition motif, LPXTG.^37,38^

Finally, flexibility in a structurally-conserved loop near the active site between the β7 and β8 strands (the β7-β8 loop) varies amongst SrtA enzymes. For example, in saSrtA, this loop shifts between two conformations in the apo and ligand-bound structures, PDB IDs 1IJA and 2KID (**Figure 1A**).^39,40^ In *Bacillus anthracis* SrtA (baSrtA), the β7-β8 loop undergoes a disordered-to-ordered transition upon ligand binding, PDB IDs 2RUI and 2KW8 (**Figure 1A**).^41,42^ In *Streptococcus pyogenes* SrtA (spySrtA), the conformation of the loop is similar in both apo and ligand-bound structures, PDB IDs 3FN5 and 7S51 (**Figure 1A**).^33,43^ The functional relevance of these differing SrtA characteristics is not well understood.

**Figure 1.**
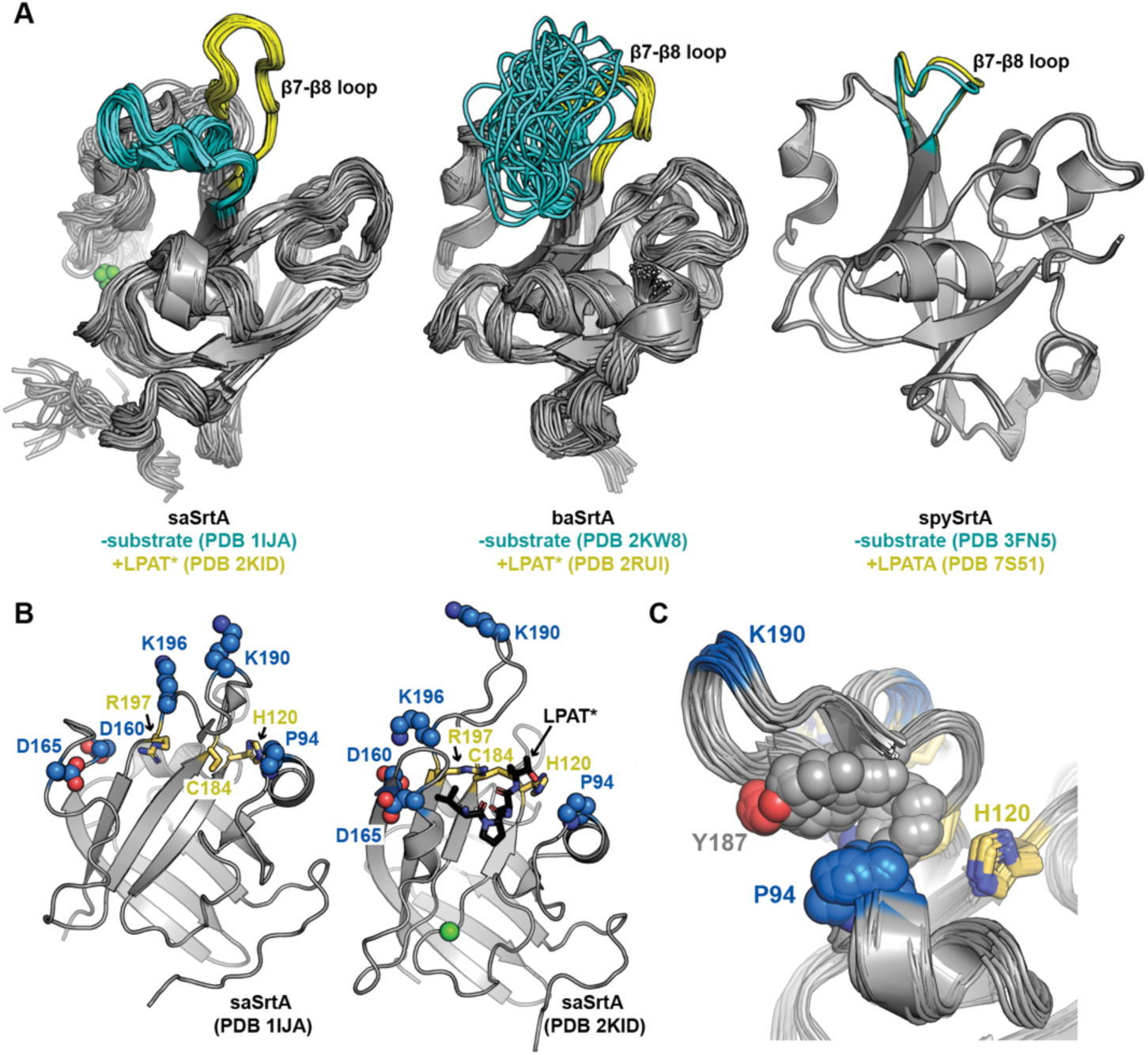
Experimental structures highlight differences in β7-β8 loop conformations without substrate bound, and an interaction between P94 and Y187 in S. aureus SrtA (saSrtA). (**A**) Experimental structures of S. aureus SrtA (saSrtA, PDB IDs 1IJA and 2KID),^39,40^ B. anthracis SrtA (baSrtA, PDB IDs 2KW8 and 2RUI),^41^ and S. pyogenes SrtA (spySrtA, PDB IDs 3FN5 and 7S51)^33,43^ are shown in cartoon representation with the β7-β8 loops colored cyan in the - substrate and yellow in the +substrate conformations. The calcium (Ca^2+^) ion is shown as a green sphere for saSrtA, which is allosterically activated by calcium. The saSrtA and baSrtA structures were determined using NMR and all available states are shown. (**B**) Mutated positions in the saSrtA5M pentamutant are highlighted as spheres and colored by heteroatom (C=blue, O=red, N=blue). The catalytic residues (H120, C184, R197) are shown as sticks and colored by heteroatom (C=yellow, N=blue, S=gold). The LPAT* peptidomimetic is shown for PDB ID 2KID as black sticks and colored by heteroatom. Ca^2+^ is shown as a green sphere. (**C**) All available NMR states are shown for saSrtA in its inactive state (PDB ID 1IJA), with the catalytic residues (H120, C184, R197) as yellow sticks and P94 and Y187 side chains as spheres. All highlighted residues are colored by heteroatom (C=yellow for catalytic residues, blue for P94, and gray for Y187, O=red, N=blue).

Directed evolution experiments previously identified a pentamutant (P94R/D160N/D165A/K190E/K196T) of saSrtA with >120-fold increased catalytic efficiency over wild-type.^44^ Subsequent work added two additional mutations (either E105K/E108A or E105K/E108Q), creating a heptamutant.^45,46^ Although the heptamutant (saSrtA7M) is approximately 3-fold less efficient than the pentamutant (saSrtA5M), it remains more active than the wild-type enzyme and does not require the Ca^2+^ cofactor; therefore, it is widely used in SML methodologies.^46,47^ As discussed in the original saSrtA5M work, the dramatic increase in catalytic efficiency is largely due to a substantial decrease in *K*_m_ with respect to recognition of the LPETG peptide substrate.^44^ Chen *et al.* reasoned that this is likely influenced by the location of the mutated residues and the resulting effects on the conformation of the structurally-conserved loops near the ligand binding groove, like the β7-β8 loop.^44^ Consistent with this, the K190E and K196T mutations are located within the β7-β8 loop (Figure 1B). Furthermore, D160N and D165A are within the β6-β7 loop, which undergoes a disordered-to-ordered transition upon calcium binding and was previously shown to affect specificity in the P4 substrate position (Figure 1B).^44,48–50^ While P94 is adjacent to the ligand binding groove, it is not located in either of these loops, although of the 5 mutated sites, it is in closest proximity to the ligand itself (Figure 1B).

In light of these observations, we wanted to better understand the saSrtA conformational transition between apo and ligand-bound structures, PDB IDs 1IJA and 2KID (Figure 1A). We observed that the side chain of P94 appears to make stabilizing hydrophobic interactions with Y187 and W194, both in the β7-β8 loop, only in the ligand-free conformation (Figure 1C). When mutated to P94R, we reasoned that these interactions would be disrupted, leading to an active conformation that mimics the ligand-bound form of the enzyme, wherein the β7-β8 loop is in the ‘open,’ upwards orientation (Figure 1A). However, we also observed that E189 in the β7-β8 loop is potentially positioned to positively interact with the Arg in the P94R saSrtA variant, which we hypothesize could favor the inactive, loop ‘down’ conformation. Conversely, the LPETG peptide substrate in the directed evolution experiment contained a P2 Glu that could be beneficial to the P94R saSrtA variant by introducing an additional enzyme-substrate interaction. Notably, P94S was reported in early rounds of the directed evolution engineering experiment, and the stereochemical consequence of Ser versus the wild-type Pro is not known.^44^ To answer these questions and better understand the effect of mutation at the P94 position, we created 18 P94X mutations, excluding only P94C, which we reasoned may form a disulfide bond with C184. We tested the relative activity of all P94X variants and wild-type saSrtA with four substrate sequences that differed at the P2 position: LP**A**TG, LP**E**TG, LP**K**TG, and LP**S**TG. We found several P94X variants that outperformed the single P94R-containing mutant. When we replaced this mutation in saSrtA5M with either P94A or P94D, we saw enhanced activity for certain LP**X**TG sequences. Consistent with this, sortase-mediated ligation experiments showed ∼2-fold greater product formation for P94D saSrtA5M as compared to saSrtA5M using a LP**K**TG substrate. As previously shown in the development of saSrtA5M, enyzme kinetics assays confirmed that these results are largely driven by a decrease in *K*_m_. Taken together, our results suggest that a thorough understanding of structure-function relationships in sortase enzymes can be used to combine rational design with high throughput techniques to develop improved tools for the broad SML protein engineering field.

## Materials and Methods

### Protein expression and purification

Recombinant sortase variant expression and purification was performed as previously described.^33,36,51–53^ The wild-type saSrtA sequence used was UniProt ID SRTA_STAA8, catalytic domain residues 60-206. Prior to the protein sequence, we included a 6x-His-tag and TEV protease cleavage site (sequence MESSHHHHHHENLYFQS), as in our previous work.^53^ Briefly, pET28a(+) plasmids (GenScript) for each variant were transformed into *Escherichia coli* BL21 (DE3) competent cells. One-liter growths using LB media and kanamycin (50 μg/mL) were grown at 37 °C to an optical density of 0.6-0.8, followed by induction with 1 mM isopropyl β-D-thiogalactoside (IPTG). The temperature was shifted to 18 °C for 18-20 hours. Cell pellets were harvested via centrifugation at 6500 rpm over 10 min. Cells were lysed with SrtA lysis buffer (0.05 M Tris pH 7.5, 0.15 M NaCl, 0.5 mM EDTA) and the lysate was sonicated for 1 minute in a 50 W, 50% duty cycle. The lysate was centrifuged for 30 minutes at 17500 rpm. The supernatant was filtered and loaded onto a Cytiva HisTrap HP (5 mL) column. The column was washed in SrtA wash buffer (0.05 M Tris pH 7.5, 0.15 M NaCl, 0.02 M imidazole, 1 mM tris(2-carboxyethyl)phosphine hydrochloride (TCEP)), followed by elution with SrtA elution buffer (0.05 M Tris pH 7.5, 0.15 M NaCl, 0.3 M imidazole, 1 mM TCEP) using a linear gradient over 20 column volumes (CV). Purified protein was concentrated using a Millipore Amicon Ultra Centrifugal Filter (10 kDa filter). In this work, we did not add TEV protease to our purified proteins, which we previously showed does not affect saSrtA activity.^51^ The concentrated protein was further purified via size exclusion chromatography (SEC) on a Cytiva HiLoad 16/600 Superdex 75 pg column. Protein aliquots were flash frozen with liquid nitrogen and stored at −80 °C. Protein identity was confirmed using SDS-PAGE and LC-ESI-MS (**Figure S1, Table S1**).

### Peptide synthesis

Peptide substrates were synthesized using the Biotage^®^ Initiatior+ Alstra^TM^ synthesizer. Synthesized peptides included the sequences: Abz-LP**X**TGGK(Dnp), with X = A, E, K, and S, Abz = 2-aminobenzoic acid, and Dnp = 2,4-dinitrophenyl. All peptides were synthesized on Fmoc Rink-Amide MBHA resin (Anaspec) on 0.1 mmol scale, resulting in peptide substrates with C-terminal primary amides (-NH_2_). Coupling reactions were performed in NMP using 5.0 molar equivalents of each appropriately protected amino acid or building block, 4.9 molar equivalents of HBTU, and 10.0 molar equivalents of DIPEA with microwave heating for 5 minutes at 75 °C. Installation of the 2,4-dinitrophenyl chromophore and Abz fluorophore was achieved through coupling of commercially available Fmoc-L-Lys(Dnp)-OH (ApexBio) or Boc-2-aminobenzoic acid (Chem-Impex International), respectively. Fmoc removal between coupling cycles was achieved using a solution of 20% piperidine in NMP. After syntheses were completed, the resin-bound peptides were washed with dichloromethane and diethyl ether. Peptides were then cleaved from the solid support using 10 mL of 95:2.5:2.5 TFA/TIPS/H_2_O for 3 h at room temperature. The cleaved peptide solutions were then concentrated using a rotary evaporator, and the remaining residue was precipitated from dry ice chilled diethyl ether. The resulting solid was collected by centrifugation and allowed to air dry overnight. Once dried, the crude peptides were dissolved in a minimum volume of acetonitrile, diluted with ∼10 mL of water and lyophilized. The crude peptides were then purified using a Dionex Ultimate 3000 HPLC system equipped with a Phenomenex Luna 5 μm C18(2) 100 Å column (10 x 250 mm) [aqueous (95% water, 5% MeCN, 0.1% formic acid) / MeCN (0.1% formic acid) mobile phase at 4.0 mL/min, method: hold 10% MeCN 0.0-2.0 min, linear gradient of 10-90% MeCN 2.0-15.0 min, hold 90% MeCN 15.0-17.0 min, linear gradient of 90-10% MeCN 17.0-17.01 min, re-equilibrate at 10% MeCN 17.01-19.0 min)]. Purified peptide fractions were concentrated on a rotary evaporator, lyophilized, and stored at −20 °C. The identity and purity of all peptides was verified by LC-ESI-MS (**Table S1**) and RP-HPLC, respectively. For use in biochemical assays, peptides were dissolved in 100% DMSO at a final concentration of ∼2-5 mM and stored at −20 °C.

### LC-ESI-MS Characterization of Sortase Variants and Peptides

Mass spectrometry characterization was achieved using a Advion CMS expression^L^ mass spectrometer interfaced with a Dionex Ultimate 3000 HPLC system. For pure peptides and reactions involving peptide substrates, separations upstream of the mass spectrometer were achieved with a Phenomenex Kinetex 2.6 μm C18 100 Å column (100 x 2.1 mm) [aqueous (95% water, 5% MeCN, 0.1% formic acid) / MeCN (0.1% formic acid) mobile phase at 0.3 mL/min, method: hold 10% MeCN 0.0-1.0 min, linear gradient of 10-90% MeCN 1.0-7.0 min, hold 90% MeCN 7.0-9.0 min, linear gradient of 90-10% MeCN 9.0-9.1 min, re-equilibrate at 10% MeCN 9.1-13.4 min]. For sortase variants, separations upstream of the mass spectrometer were achieved with a Phenomenex Aeris^TM^ 3.6 μm WIDEPORE C4 200 Å column (100 x 2.1 mm) [aqueous (95% water, 5% MeCN, 0.1% formic acid) / MeCN (0.1% formic acid) mobile phase at 0.3 mL/min, method: hold 10% MeCN 0.0-1.0 min, linear gradient of 10-90% MeCN 1.0-7.0 min, hold 90% MeCN 7.0-10.0 min, linear gradient of 90-10% MeCN 10.0-10.1 min, re-equilibrate at 10% MeCN 10.1-13.25 min]. Deconvolution of protein charge ladders was achieved using Advion Data Express (version: 6.9.43.1) software.

### Sortase activity assays

Enzymatic activity was tested using fluorescence assays with the BioTek Synergy H1 plate reader. The following concentrations were used, unless otherwise noted in the text: 1 μM enzyme, 0.05 mM peptide, 5 mM hydroxylamine (H_2_NOH). Assays were performed using sortase running buffer (0.05 M Tris pH 7.5, 0.15 M NaCl, 0.01 M CaCl_2_). Assays were run for 2 h at room temperature using an excitation wavelength of λ=320 nm. Data was recorded at the emission wavelength, λ=420 nm. Time-matched fluorescence of the peptide alone (negative control) was subtracted from each data point. All assays were run as independent replicates with N=3 for each enzyme-substrate pair, and standard deviation errors were calculated. Calculations and graphing were done using Excel and GraphPad Prism.

To complement fluorescence assays, as well as to monitor product formation from sortase-mediated ligation reactions, select pairings of sortase enzyme and peptide substrates were also monitored by HPLC. This was achieved using a Dionex Ultimate 3000 HPLC system and a Phenomenex Kinetex 2.6 μm C18 100 Å column (100 x 2.1 mm) [aqueous (95% water, 5% MeCN, 0.1% formic acid) / MeCN (0.1% formic acid) mobile phase at 0.3 mL/min, method: hold 10% MeCN 0.0-1.0 min, linear gradient of 10-90% MeCN 1.0-7.0 min, hold 90% MeCN 7.0-9.0 min, linear gradient of 90-10% MeCN 9.0-9.1 min, re-equilibrate at 10% MeCN 9.1-13.4 min]. The following reagent concentrations were used: 0.1 μM enzyme, 0.05 mM peptide, and 5 mM hydroxylamine (H_2_NOH) or 5 mM diglycine (Gly-Gly) in sortase running buffer (0.05 M Tris pH 7.5, 0.15 M NaCl, 0.01 M CaCl_2_). Assays were run for 2 h at room temperature and monitored at 360 nm. The extent of product formation was estimated from peak areas for the unreacted peptide substrate (Abz-LP**X**TGGK(Dnp)) and the C-terminal cleavage fragment (GGK(Dnp)). The identity of all relevant peptide peaks was confirmed using LC-ESI-MS (**Table S1**), as described above.

### Enzyme kinetics assays

Reaction mixtures were prepared in a total volume of 200 µL and included the following buffer and nucleophile components: 0.05 M Tris pH 7.5, 0.15 M NaCl, 0.01 M CaCl_2_, 1 mM hydroxylamine (H_2_NOH), and a final DMSO concentration of 5% (*v/v*). For all assays, the enzyme concentration was 2.79 µM. The substrate used was Abz-LP**E**TGGK(Dnp), at variable concentrations, which included 10 µM, 25 µM, 50 µM, 100 µM, 250 µM, 500 µM, and 1 mM. Assays were run in triplicate. Reaction time points (10 s, 60 s, 120 s, and 180 s) were obtained by quenching aliquots of each reaction with glacial acetic acid (7:1 final ratio of reaction mixture to acetic acid). Samples were analyzed via HPLC using a Dionex Ultimate 3000 HPLC system and a Phenomenex Kinetex 2.6 μm C18 100 Å column (100 x 2.1 mm) [aqueous (95% water, 5% MeCN, 0.1% formic acid) / MeCN (0.1% formic acid) mobile phase at 0.3 mL/min, method: hold 10% MeCN 0.0-1.0 min, linear gradient of 10-90% MeCN 1.0-7.0 min, hold 90% MeCN 7.0-9.0 min, linear gradient of 90-10% MeCN 9.0-9.1 min, re-equilibrate at 10% MeCN 9.1-13.4 min]. Samples were monitored at 360 nm, and two major peaks were observed for the unreacted peptide substrate (Abz-LP**E**TGGK(Dnp)) and the C-terminal cleavage fragment (GGK(Dnp)). The identity of these peaks was confirmed using LC-ESI-MS (as described above), and initial velocities (µM/sec) were estimated from their corresponding peak areas. GraphPad Prism was used to fit the Michaelis-Menten equation, and extract V_max_ and *K*_m_ values for each replicate; *k*_cat_ was calculated using the equation V_max_= *k*_cat_*[Enzyme]_total_.

### Structural Modeling

The AlphaFold3 server was used for all structural modeling, including of P94X saSrtA variants with and without Ca^2+^, or with a peptide substrate sequence (either LP**X**TG or AQALP**X**TG as indicated in the Supplemental Information) and Ca^2+^.^54^ The N-terminal AQA sequence was derived from the endogenous saSrtA target, Spa (UniProt ID SPA_STAA8), and was necessary for proper substrate binding in the wild-type sequence and for modeling with selected variants and LP**A**TG. PyMOL was used for structural figures and analyses, including the APBS plug-in to calculate electrostatic surface potential maps.

## Results

### Relative activities of P94X saSrtA variants with the LPETG substrate

In order to assess the impact of variations in the P94 saSrtA position, we first expressed and purified 18 single mutations. We included all amino acids except for the P94C variant due to the proximity of this residue to the catalytic C184 position, with an average distance of 8.3 Å (minimum distance =7.7 Å) between the Cα atoms of P94 and C184 over the 25 NMR states of PDB 1IJA. Although this is outside the range of 3.0-7.5 Å previously observed for disulfide bonded cysteines, we reasoned that this was close enough that potential disulfide bond formation may perturb overall folding of the enzyme.^55^ All variants were expressed and purified using similar protocols as described previously and as in the Materials and Methods (**Figure S1, Table S1**).^33,36,51–53^

To compare relative activities of our P94X saSrtA variants, we utilized a fluorescence resonance energy transfer (FRET)-based peptide cleavage assay. We labeled a LPXTG-containing peptide at the N-terminus with a 2-aminobenzoyl (Abz) fluorophore, and at the C-terminus with a 2,4-dinitrophenyl (Dnp) quencher, as previously described and as in the Materials and Methods.^33,36,37,51–53,56^ The general sequence of our synthesized peptide substrates was Abz-LP**X**TGGK(Dnp), with X=Glu here, as this was the substrate used in the directed evolution study that produced the saSrtA pentamutant (saSrtA5M).^44^ For clarity, we will omit the Abz- and GK(Dnp) residues when referring to these substrates moving forward, e.g., LP**E**TG for Abz-LP**E**TGGK(Dnp). Upon sortase-mediated cleavage between the P1/P1’, or T/G, positions, we observed an increase in Abz fluorescence. While this assay only monitors the first step of the reaction, recognition of the substrate and formation of the acyl-enzyme intermediate was previously shown to be the rate-limiting step of the overall sortase reaction.^57–59^ Furthermore, a previous kinetic analysis of P94S saSrtA revealed a 3-fold decrease in *K*_m_ for this mutant with no change in *k*_cat_ (WT: *K*_m_ = 7.6 mM, *k*_cat_=1.5 s^-1^; P94S: *K*_m_ = 2.5 mM, *k*_cat_=1.6 s^-1^, for the LP**E**TG substrate), suggesting that variation at this residue affects this initial step of the reaction.^44^

In our first experiment, we tested 19 saSrtA enzymes, including our variants and wild-type (WT), with the LP**E**TG peptide (Figures 2A**, S2**). Initially, we used an enzyme concentration of 5 μM and the peptide substrate at 50 μM, consistent with our previous work.^33,36,37,51–53,56^ However, this proved to be too high of a concentration to capture the early timepoints for several of our variants, and we decreased our P94X saSrtA concentration to 1 μM, as described in the Materials and Methods. In order to directly compare relative activities with the wild-type enzyme, we decided to focus our analyses on a time point relatively early in our t=2 h experiment, before we observed any fluorescence plateaus. Therefore, we chose to normalize relative fluorescence at t=20 min as compared to the WT saSrtA enzyme (Figure 2B). Notably, our results indicated that only the P94F variant showed lower activity than the WT enzyme. Interestingly, while P94R (the mutation included in saSrtA5M) had 1.89-fold higher relative fluorescence at t=20 min, as compared to WT, this was far from the highest activity variant. Indeed, several mutations resulted in >2-fold higher changes as compared to WT, including A (3.0-fold), D (2.6), E (2.3), G (3.4), H (2.9), K (2.1), N (2.4), Q (2.5), S (3.0), and T (3.3) (Figure 2B). Of these, P94G, P94S, and P94T showed ≥3-fold increases in relative fluorescence on average as compared to WT (Figure 2B). Of the additional positively-charged amino acids, P94K (2.1-fold as compared to WT) behaved similarly to P94R, while P94H was relatively more active at t=20 min (2.86-fold as compared to WT) (Figure 2B).

**Figure 2.**
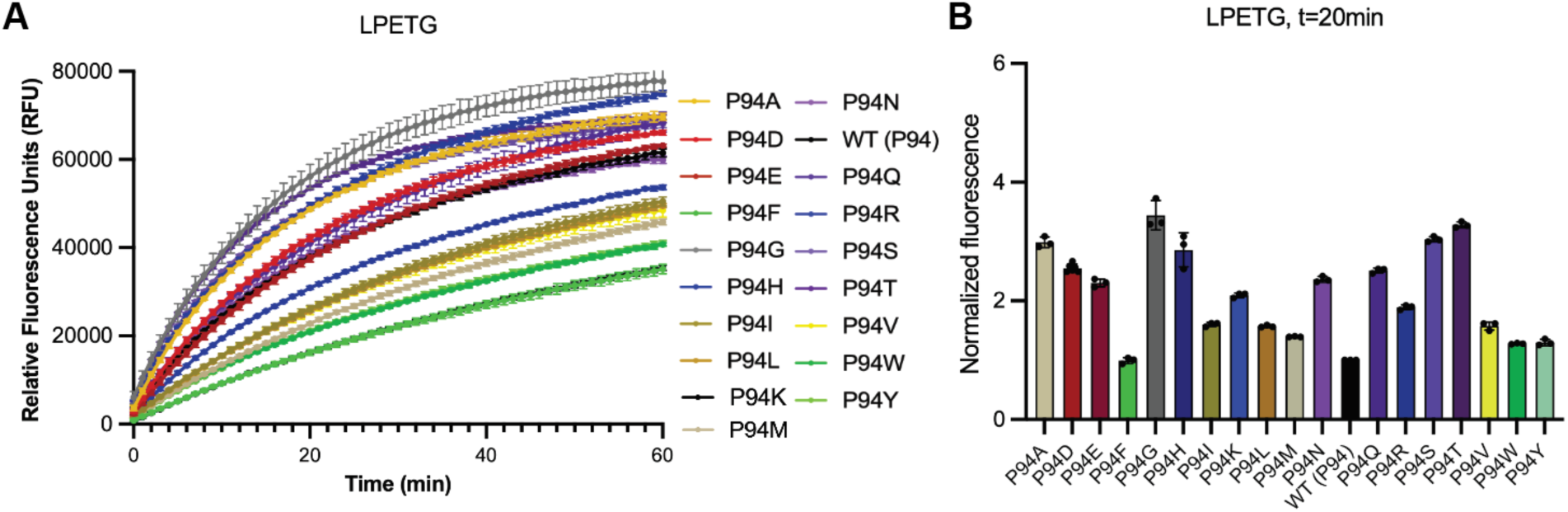
Activity assay data for P94X saSrtA variants and the LPETG substrate sequence. (**A**) Triplicate data (shown as averaged values with error bars equal to standard deviation) are shown for the P94X saSrtA variants during the first 60 min of the activity assay (total time = 2 h). Variants are colored as in the key, and correspond to the colors used in (**B**). The full 2 h time course data is in **Figure S2**. Calculated initial velocities are in **Figures S7-S8** and **Table S4**. (**B**) Normalized fluorescence (as averaged values ± standard deviation) for the P94X saSrtA variant data at the t=20 min time point. Normalization is as compared to the wildtype (WT) data, which is equal to 1.

To begin to understand the effects of our panel of P94 mutations, we used AlphaFold3 to see if structural models showed a difference in the position of the loop that contains this mutation.^54^ The P94 position is located on an α-helix between the β2 and β3 strands, and is situated almost directly across the peptide-binding cleft from the catalytic C184 residue (Figure 3A). Based on experimental structures, P94 forms a hydrophobic interaction with Y187 in the ligand-free state (PDB 1IJA), a position in the β7-β8 loop (Figure 1C).^39^ As discussed above, there is a conformational change in the β7-β8 loop upon substrate-binding that dramatically increases this distance (Figure 3B). In the NMR experimental structures, the loop ‘closed, down’ inactive conformation revealed a distance of 7.3 ± 0.2 Å (PDB 1IJA), and the loop ‘up, open’ active conformation was 17.1 ± 0.3 Å (PDB 2KID) (Figures 1A**, 3B**). Therefore, we predicted that a P94X mutation may affect overall flexibility and dynamics in the β7-β8 loop. As discussed in detail in the Supplemental Information, we used AlphaFold3 to model all P94X saSrtA variants ± Ca^2+^, as well as several with the LPETG peptide substrate (**Figures S3-6, Tables S2-3**). Ultimately, we did not observe widespread differences in the P94X-Y187 distance based on mutation at P94X, although it would be interesting to use experimental methods to better interrogate this system.

**Figure 3.**
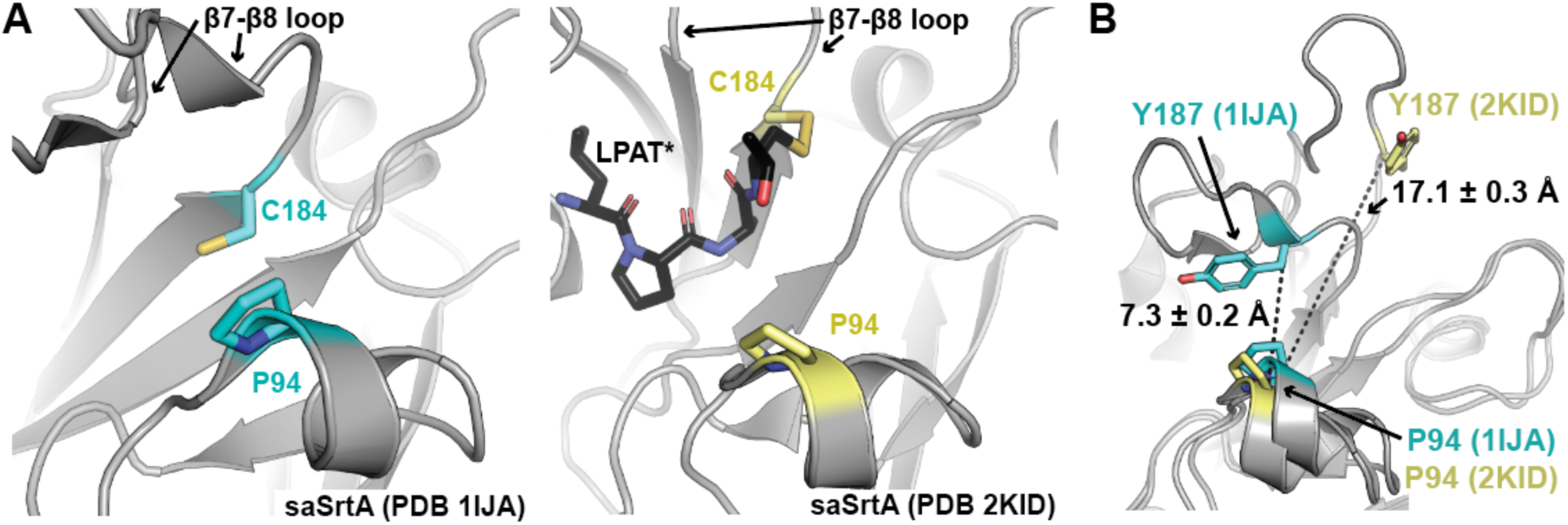
Distance between P94 and Y187 in saSrtA experimental structures. **(A)** The location of P94 with respect to the catalytic C184 residue (or Y187 in **B**) is highlighted in the wild-type saSrtA structures (PDB ID 1IJA (left) and 2KID (right).^39,40^ SaSrtA is in gray cartoon, and the side chain atoms of P94 and C184 are shown as sticks and colored by heteroatom (C=cyan for 1IJA and yellow for 2KID, N=blue, S=gold). The LPAT* peptidomimetic in PDB 2KID is in black sticks and colored by heteroatom (C=black, N=blue, O=red, S=gold). In (**B**), distances between Cα atoms are labeled and shown as black dashes.

### Relative activities of P94X saSrtA variants with LPATG, LPKTG, and LPSTG

We next wanted to directly test the effect of P2 (LP**X**TG) identity on relative activity. The P94 amino acid is in a stereochemical environment such that we predicted it may be able to directly interact with the P2 (LP**X**TG) position of the substrate. Although our AlphaFold3 models containing peptide did not show any specific interactions, we wondered if there may be transient interactions during substrate binding/unbinding that could differentially affect the *k*_on_ and/or *k*_off_ for different substrate sequences. While this would be challenging to experimentally test due to the expected relatively high overall *K*_m_ for SrtA and peptide substrates (in the mM range), we reasoned that we could estimate the effects on these parameters by measuring relative activities. Thus, we tested our P94X variants with sequences that varied at the P2 position. In addition to LP**E**TG, we chose to test P2=A, K, and S due to the differing chemical properties of these amino acids. For example, while E is negative, K is positive, S is polar and uncharged, and A is small and hydrophobic.

Peptide substrates were synthesized as described above and as in the Materials and Methods, and activity assays were performed as with the LP**E**TG substrate (Figure 2). Again, we focused our analyses on the t=20 min timepoint (Figures 4A-C). However, we also calculated initial velocities (in RFU/min) for all P94X variants with all substrates, including LP**E**TG (**Figure S7-S8, Table S4**). Our data revealed that while there were particular P94X variants that showed the highest relative activities for several of these substrates, specific P2 preferences also emerged (Figures 4A-C). For example, if we look at variants with an average increase in relative activity of >3-fold as compared to WT, they were (with shared P94X variants underlined): LP**A**TG: A, D, E, <u>G</u>, <u>S</u>, and <u>T</u>; LP**E**TG: <u>G</u>, <u>S</u>, and <u>T</u>; LP**K**TG: D, E, <u>G</u>, <u>S</u>, and <u>T</u>; and LP**S**TG: A, D, E, <u>G</u>, N, <u>S</u>, and <u>T</u>, (Figures 2**, 4**).

**Figure 4.**
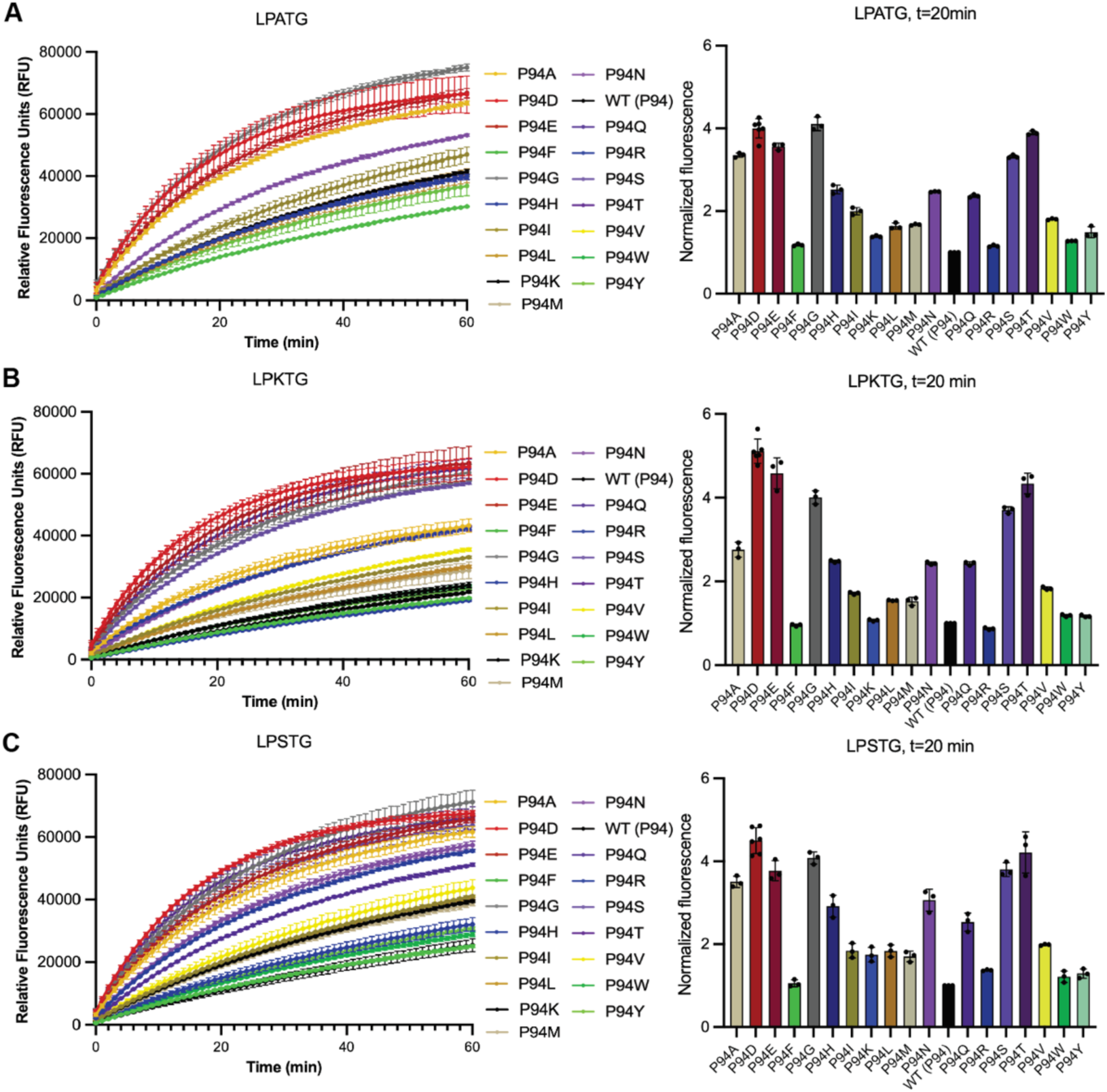
Activity assay data for P94X saSrtA variants and LP<u>A</u>TG, LP<u>K</u>TG, and LP<u>S</u>TG substrate sequences. Triplicate data (shown as averaged values with error bars equal to standard deviation) are shown for the P94X saSrtA variants with LP**A**TG (**A**), LP**K**TG (**B**), and LP**S**TG (**C**) peptide substrates during the first 60 min of the activity assay (total time = 2 h). Variants are colored as in the key, and corresponding to the colors used in Figure 2. The 2 h time course data is in **Figure S2**. Calculated initial velocities are in **Figures S7-S8** and **Table S4**.

To better investigate specific positive and/or negative preferences, we created a heat map of the average normalized fluorescence values for each P94X variant as compared to its relative fluorescence for LP**E**TG at t=20 min (Figure 5A). In general, the lowest relative activities are for LP**K**TG, which we reasoned may be due to the proximity of R197, and repulsive electrostatic interactions. We also analyzed specificity for P2 identify in the WT saSrtA enzyme (Figure 5B), and saw that at t=20 min, WT saSrtA activity, as compared to LP**E**TG, is: 0.7x for LP**A**TG, 0.6x for LP**K**TG, and 0.7x for LP**S**TG. Therefore, the wild-type enzyme prefers E > A > S > K at the P2 position (Figure 5B). Our results are largely consistent with previous data that investigated P2 specificity in saSrtA, which shows relatively minor variability in a ligation assay (t=30 min) for P2=A, E, and S (K was not tested).^60^ More dramatic differences from this previous work suggested positive preferences for P2=M (>2-fold higher activity as compared to a P2 E) and negative preferences for P2=G, I, T, and V (>2-fold lower activity as compared to a P2 E).^60^

**Figure 5.**
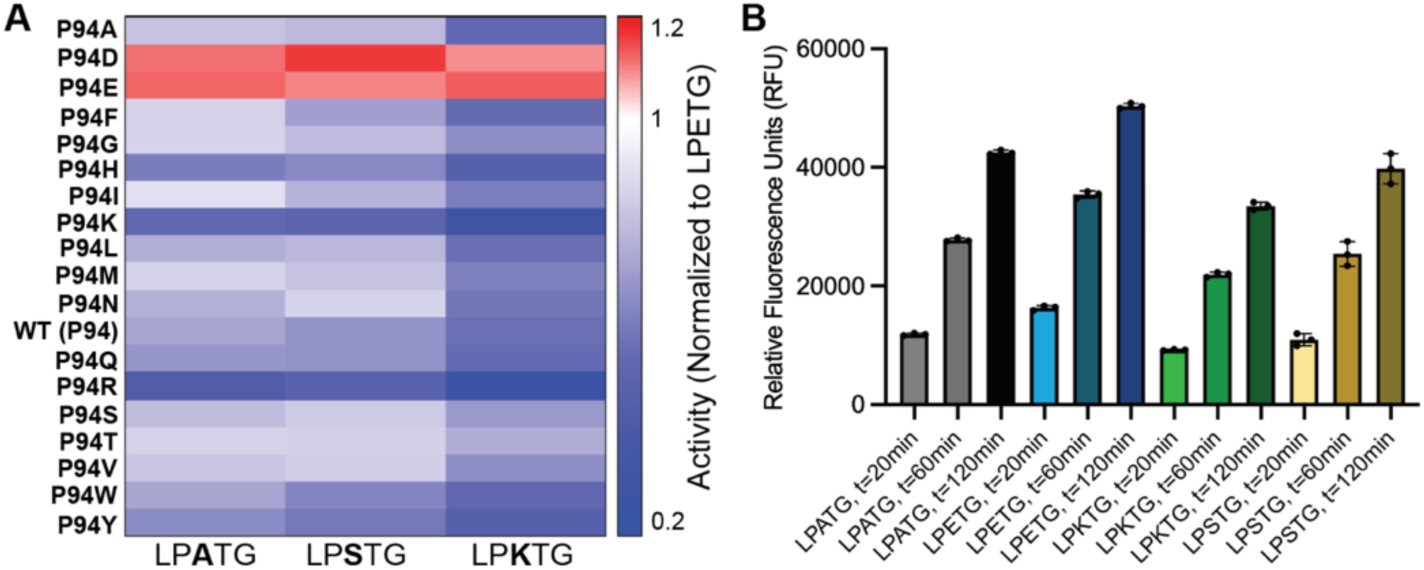
Relative P2 preferences for P94X saSrtA variants and wild-type. (A) A heat map comparing differences in relative fluorescence unit (RFU) values at t=20 min for each P94X saSrtA variant, as compared to its value for the LP**E**TG substrate (Figure 2). For example, the red colors for P94D and P94E saSrtA with the LP**A**TG, LP**K**TG, and LP**S**TG substrates indicate that these are preferred over LP**E**TG (RFU_LP**X**TG_/RFU_LP**E**TG_ is >1). (**B**) RFU values for wild-type saSrtA at t=20,60,120 min as a function of the P2 position in the LP**X**TG substrates.

As shown in our heat map analyses, there were clear preferences that were not apparent when we normalized our fluorescence data to the wild-type enzyme (Figure 5A). Most were relatively minor and in general all P94X variants (except for P94D and P9E) disfavored LP**K**TG. There was an additional P2-specific effect observed in the P94R and P94K saSrtA variant data, where, as compared to LP**E**TG, P94R average fluorescence was 0.4x for LP**A**TG, 0.3x for LP**K**TG, and 0.5x for LP**S**TG (Figure 5A). We observed similar results when we graphed the fluorescence data for select individual P94X variants and WT and all substrates (Figure 6), as well as the average relative fluorescence values for each variant/peptide pair at t=20 min (Figure 7). In general, all variants with only two exceptions (P94D and P94E) preferred the P2 Glu (LP**E**TG), followed by LP**A**TG or LP**S**TG (values within 85% of each other), then LP**K**TG (Figure 7). For P94D and P94E saSrtA enzymes, there was little difference in activity between substrates (Figures 6-7), although the P2 Glu was disfavored by both according to our heat map using t=20 min data (Figure 5A).

**Figure 6.**
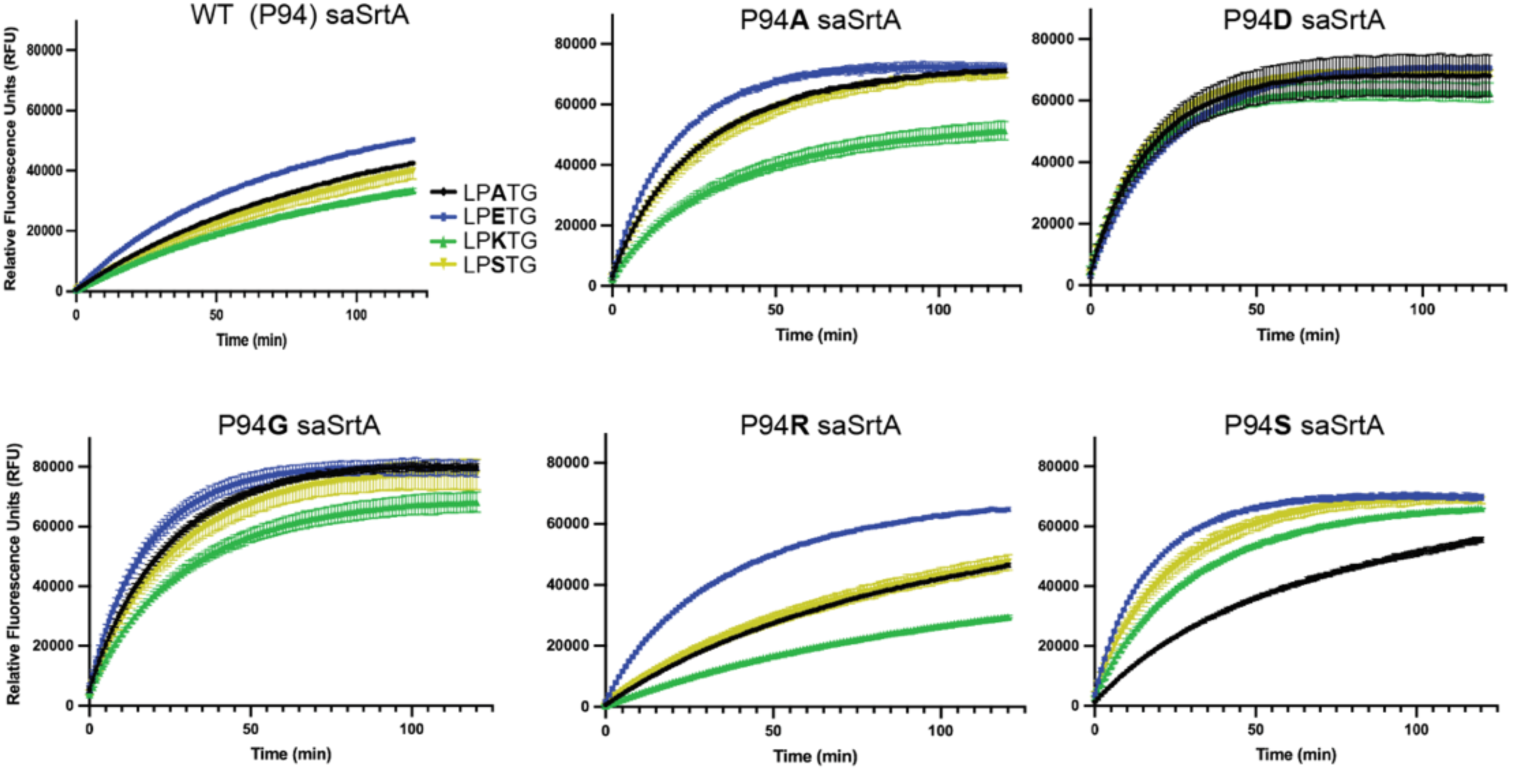
Activity assays for select variants as a function of P2 position in LPXTG substrates. Activity assays (t=120 min) for select wild-type (WT) and P94X saSrtA variants as a function of the P2 position in the LP**X**TG substrates. Curves are colored as in the key to the right of the WT saSrtA data.

**Figure 7.**
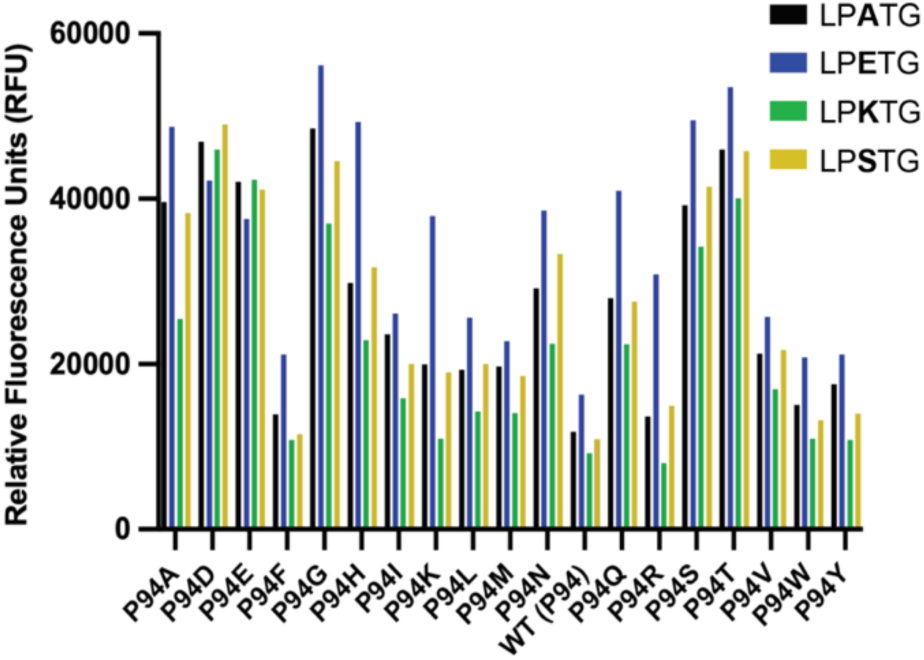
Relative fluorescence values for all P94X saSrtA assays at t=20 min. Averaged RFU values for all P94X variants with all LP**X**TG substrates at t=20 min. All experiments were performed in triplicate.

We modeled P94D, P94E, and P94R saSrtA with a LP**A**TG peptide sequence and Ca^2+^ using AlphaFold3. In all cases, initial modeling failed to properly position the substrate; therefore, we added N-terminal AQA residues and used the sequence AQALP**A**TGG to facilitate the modeling (**Figure S4**). The electrostatic potential surface maps of the modeled saSrtA variants revealed that the relative charge does change with the positively charged P94R versus negatively charged P94D or P94E residues (Figure 8). These analyses also confirmed that the opposite face of the substrate binding pocket in this region is positively charged, due to the R197 residue (Figure 8). Our previous work revealed that R197 (R216 in *Streptococcus pyogenes* SrtA, PDB ID 7S51) stabilizes the backbone of the substrate via hydrogen bonds.^33^ We also saw non-covalent interactions in our saSrtA-LP**E**TG models, with the R197-P3 Pro interaction positioning the guanidino group of the R197 side chain in relatively close proximity to the carboxylic acid of the P2 Glu (**Figure S3A**). This may explain why the repulsive negative preference of P94D and P94E saSrtA for the P2 Glu is less pronounced than that for P94R and a P2 Lys, because P94D/E would be additionally affected by a favorable interaction with the positively charged R197 amino acid.

**Figure 8.**
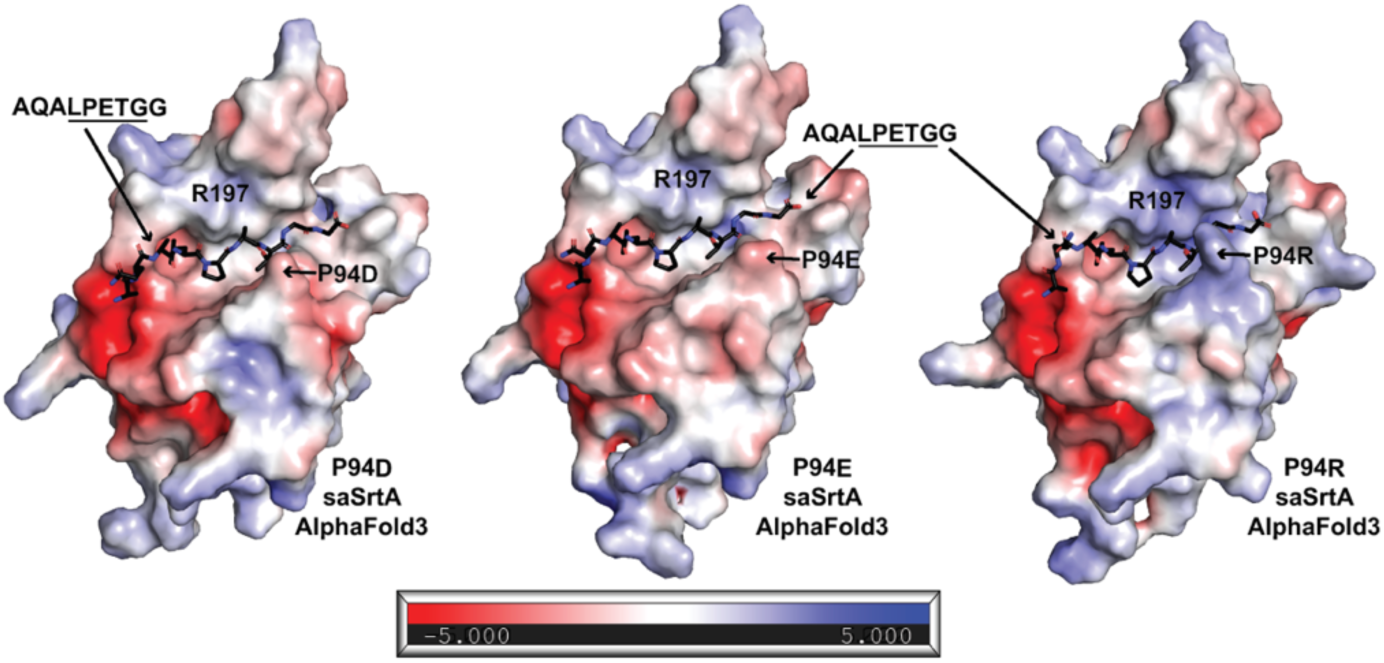
Electrostatic potential surface maps of P94D, P94E, and P94R saSrtA-substrate AlphaFold3 models. Electrostatic potential surface maps (calculated using APBS in PyMOL) for AlphaFold3 models of P94D saSrtA-AQA<u>LPETG</u>G (left), P94E saSrtA-AQA<u>LPETG</u>G (middle), and P94R saSrtA-AQA<u>LPETG</u>G (right), as labeled. The peptide substrate is shown as sticks and colored by heteroatom (C=black, O=red, N=blue). AlphaFold3 output data is in **Table S2** and **Figures S5-S6**.

### Effect of P94X mutation on saSrtA5M activities

Based on our results, we hypothesized that P94R mutation in saSrtA5M may exhibit different effects on activity for certain substrate sequences. We chose to test P94A and P94D in the context of the saSrtA5M enzyme, as these variants were relatively active compared to wild-type, and represent different chemical properties (Figures 2**, 4A-C**). We expressed and purified these variants as described in the Materials and Methods, and used our peptide cleavage assay to test their relative activities. We used enzyme concentrations of 1 μM and 0.1 μM. While the 1 μM concentrations allowed us to directly compare with our other assays, the 0.1 μM concentrations were required to capture initial activity.

Consistent with our data and previous work, the saSrtA5M variants were *used was 1 μM.* generally more active than all single P94X mutations at t=20 min, except for saSrtA5M with LP**K**TG (Figure 9). Although the P94A and P94D saSrtA5M variants tested revealed similar activities for LP**E**TG, this was not the case for the other P2 substrates. While the effect was most dramatic for LP**K**TG, as hypothesized due to the negative preference for P94R saSrtA for the P2 Lys, we found that P94A and P94D saSrtA5M also outperformed saSrtA5M (with the P94R mutation) at early time points for P2=Ala, Lys, and Ser substrates (Figure 10). In our 1 μM assays, the relative fluorescence values were approximately equal by t=60 min; however, this was not the case in our 0.1 μM assays at the t=2 hr time point (Figures 9-10). In these lower concentration assays, we again saw clear differences at the 20 min timepoint (**Figure 10B**). In addition, initial velocity measurements revealed even more dramatic effects, with initial average velocities for P94A saSrtA and P94D saSrtA as compared to saSrtA5M equal to: 1.4x and 1.6x for LP**A**TG, 1.1x and 0.9x for LP**E**TG, 2.1x and 3.3x for LP**K**TG, and 1.4x and 1.5x for LP**S**TG, respectively (**Figure 10C**). Overall, this data suggests that P2 specificity plays a role in the widely used engineered saSrtA5M enzyme, and that different P94X mutations in the saSrtA5M context may be beneficial for substrates that are not LP**E**TG.

**Figure 9.**
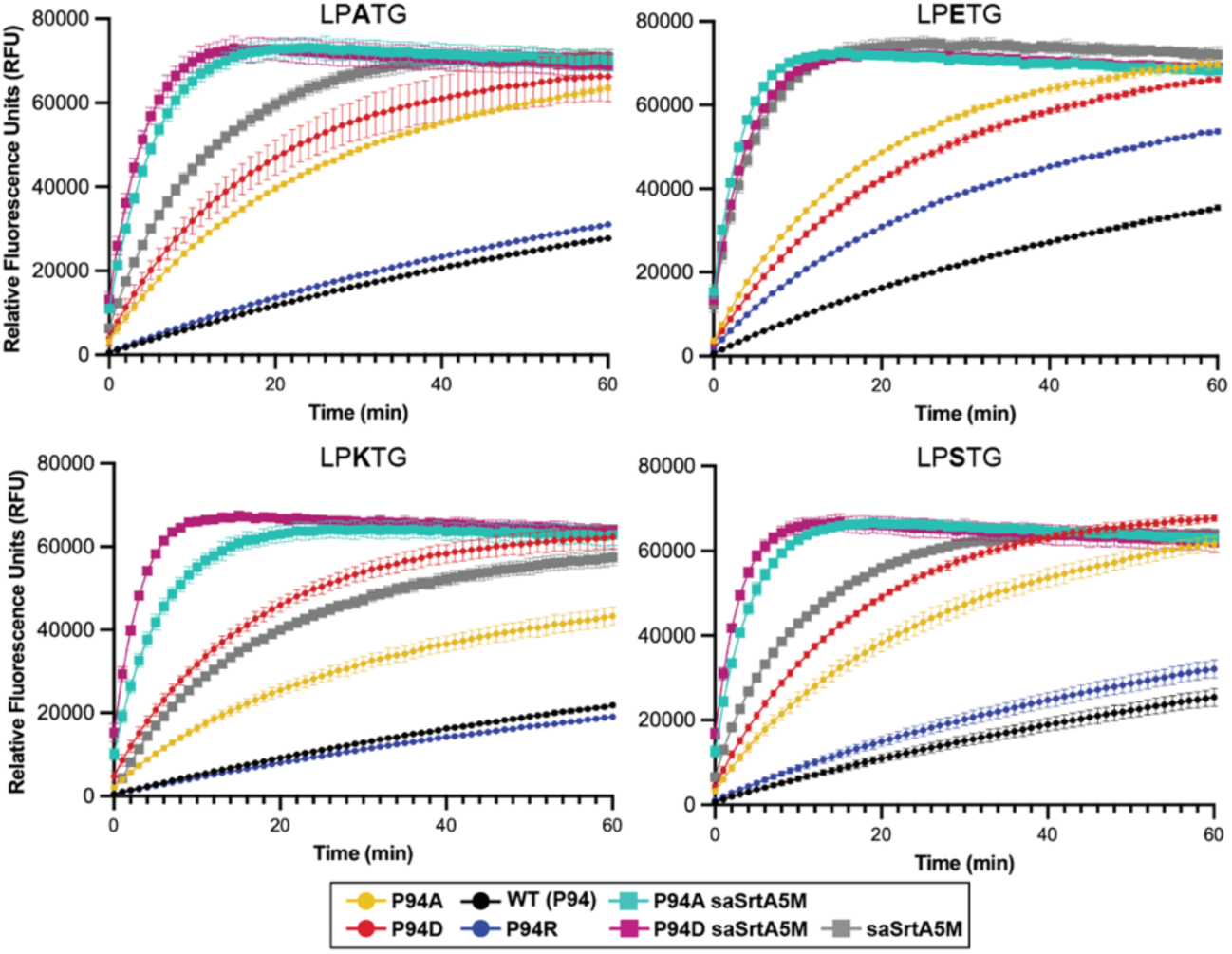
Activity assay data for select P94X saSrtA variants as compared to saSrtA5M variants. Triplicate data (shown as averaged values with error bars equal to standard deviation) are shown for select P94X saSrtA variants as compared to saSrtA5M, P94A saSrtA5M, and P94D saSrtA5M (key at bottom). Peptide substrates tested included: LP**A**TG, LP**E**TG, LP**K**TG, and LP**S**TG, as labeled. For all, the enzyme concentration

**Figure 10.**
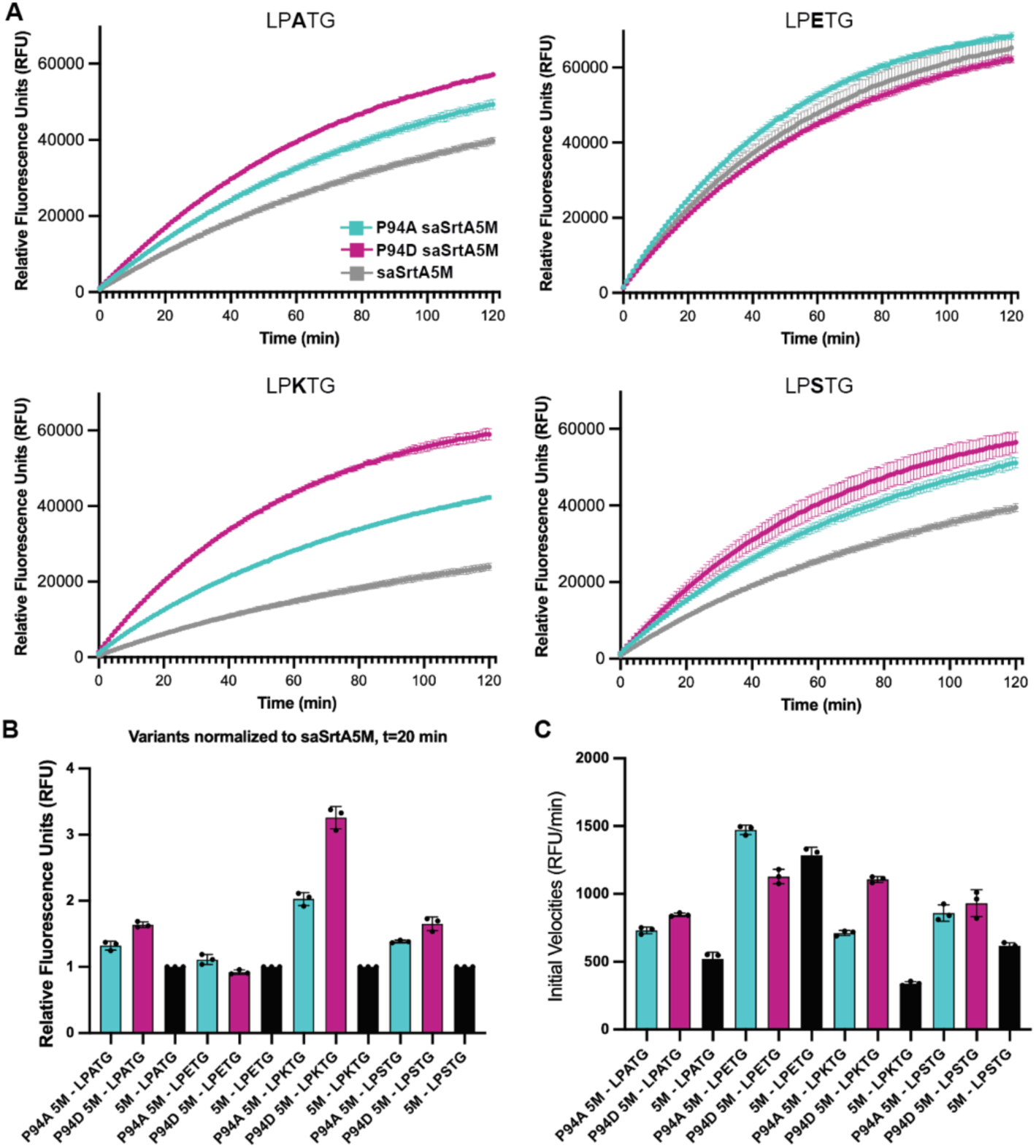
Effects of P94X mutation in the saSrtA5M pentamutant enzyme. (A) Triplicate data (shown as averaged values with error bars equal to standard deviation) for saSrtA5M, P94A saSrtA5M, and P94D saSrtA5M variants (key in top left). Peptide substrates tested included LP**A**TG, LP**E**TG, LP**K**TG, and LP**S**TG, as labeled. For all, the enzyme concentration used was 0.1 μM. (**B**) Relative fluorescence unit (RFU) values at t=20 min, as normalized to the saSrtA5M (5M) data for each peptide substrate. Therefore, saSrtA5M = 1 for each of LP**A**TG, LP**E**TG, LP**K**TG, and LP**S**TG peptides. (**C**) Initial velocities (in RFU/min) for all saSrtA5M variants tested, with each peptide substrate, shown as average values ± standard deviation. Curves and calculated values are in **Figure S7** and **Table S4**.

### Enzyme kinetics and HPLC cleavage assays using P94D saSrtA variants and saSrtA5M

To further characterize the effects of P94X mutations, we next determined the kinetic parameters (*k*_cat_ and *K*_m_) of our P94D and P94R saSrtA and saSrtA5M variants and compared them to the standard wild-type saSrtA and saSrtA5M enzymes (**Table 1**). Initial reaction rates were calculated using an HPLC assay with fixed concentrations of enzyme and the H_2_NOH nucleophile, and various Abz-LP**E**TGGK(Dnp) substrate concentrations, as previously described and in the Materials and Methods (**Figure 11a**).^61^ Importantly, our kinetics analyses were consistent with our peptide cleavage assay, as results were comparable to the relative fluorescence data described above. With respect to the literature, previously published *k*_cat_ and *K*_m_ values for sortase enzymes differ substantially, which we attribute to variations in the substrate peptide and nucleophiles used.^44,59,61–63^ Of these previous reports, the substrate/nucleophile pair most similar to our system is Abz-LPETGG-Dap(DNP)-NH_2_ with a 2 mM pentaglycine (GGGGG) nucleophile, which resulted in *k*_cat_ = 1.10 ± 0.06 s^-1^ and *K*_m_ = 8.76 ± 0.78 mM for wild-type saSrtA and *k*_cat_/*K*_m_ = 125 ± 18 M^-1^ s^-1^.^63^ While our values are lower than these (*k*_cat_ = 0.022 ± 0.001 s^-1^ and *K*_m_ = 0.185 ± 0.01 mM for wild-type saSrtA), they are shifted by a consistent factor of ∼50x (**Table 1**). As a result, our *k*_cat_/*K*_m_ = 120 ± 1 M^-1^ s^-1^ value strongly agrees with that previously reported.

**Figure 11.**
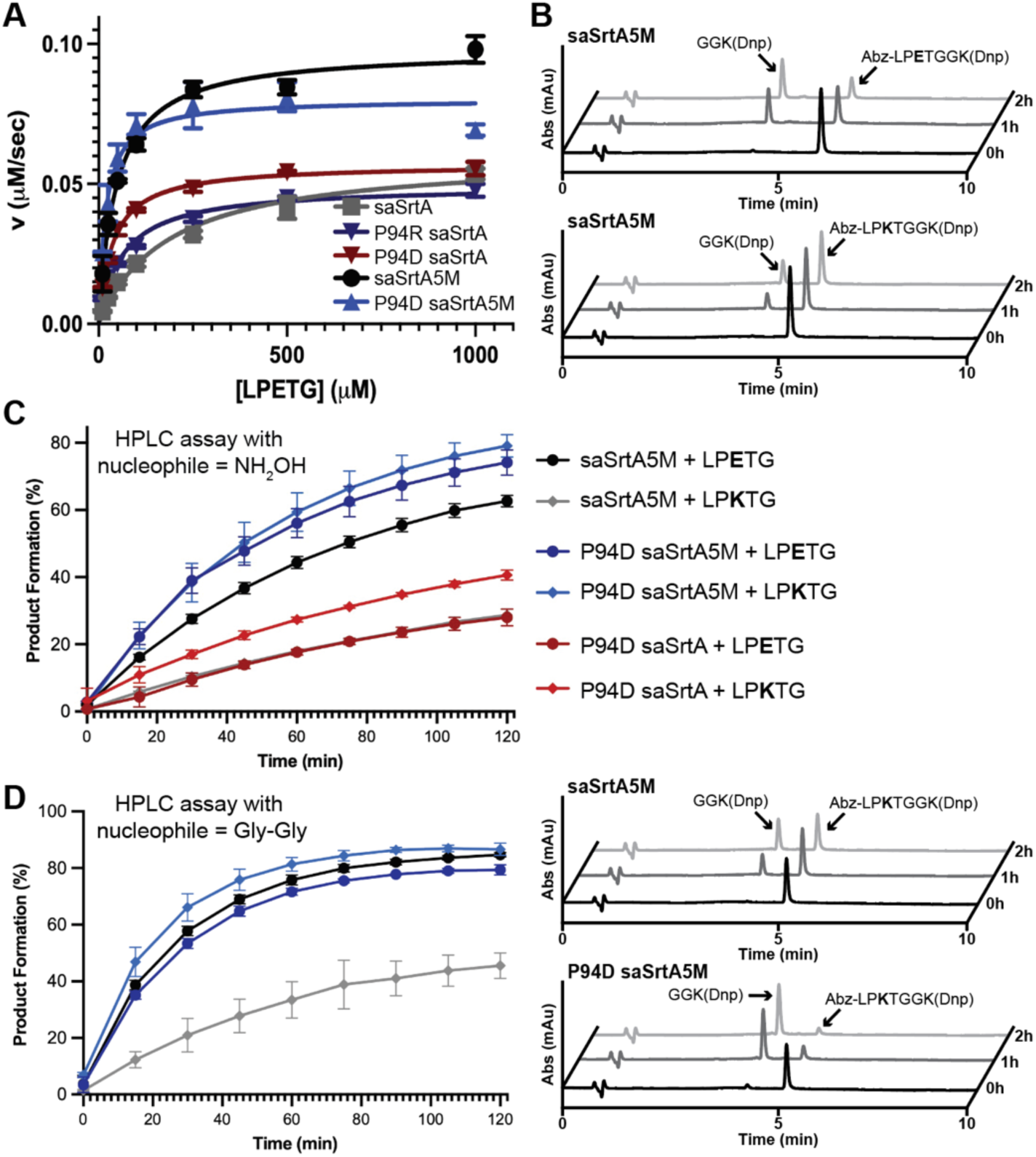
Enzyme kinetics and HPLC assays for saSrtA and saSrtA5M variants. (**A**) Average of replicate data (N=3) for enzyme kinetics assays of saSrtA variants. Data was fit using the Michaelis-Menten equation in GraphPad Prism. Individual replicate data is in **Figure S9**. (**B**) Representative HPLC traces (360 nm) of sortase-mediated reactions utilizing a H_2_NOH nucleophile. Product formation catalyzed by the saSrtA5M enzyme was higher with the LP**E**TG substrate, as compared to LP**K**TG, based on the areas of the substrate (Abz-LP**X**TGGK(Dnp)) peak and cleavage product peak (GGK(Dnp)) at indicated time points. (**C**) Quantification of sortase-mediated reactions (H_2_NOH nucleophile) using HPLC with standard deviation error bars shown. All experiments were conducted as technical triplicates and are described in the Materials and Methods. The extent of product formation was estimated from peak areas for the unreacted peptide substrate (Abz-LP**X**TGGK(Dnp)) and the C-terminal cleavage fragment (GGK(Dnp)). (**D**) Quantification (all) and representative HPLC traces for sortase-mediated ligation reactions (Gly-Gly nucleophile) using the substrates LP**E**TG or LP**K**TG. The figure legend is the same as in (**C**), with saSrtA5M in black circles (+LP**E**TG) and gray diamonds (+LP**K**TG), and P94D saSrtA5M in dark blue circles (+LP**E**TG) and blue diamonds (+LP**K**TG). HPLC traces (360 nm) are rendered as in (**B**), highlighting the enhancement in reaction conversion for the LP**K**TG substrate when using the P94D saSrtA5M enzyme. Additional representative HPLC traces for saSrtA5M, P94D saSrtA5M, and P94D saSrtA catalyzed reactions are in **Figure S10**.

**Table 1.** Kinetic characterization of saSrtA variants.

| | $k_{\text{cat}}, \text{s}^{-1}$ | $K_{\text{m}} \text{LPETG}, \mu\text{M}$ | $k_{\text{cat}}/K_{\text{m}} \text{LPETG}, \text{M}^{-1} \text{s}^{-1}$ |
| --- | --- | --- | --- |
| **WT saSrtA** | **$0.022 \pm 0.001$** | **$185 \pm 10$** | **$120 \pm 1$** |
| **P94D saSrtA** | **$0.020 \pm 0.01$** | **$37 \pm 3$** | **$550 \pm 30$** |
| **P94R saSrtA** | **$0.017 \pm 0.02$** | **$65 \pm 5$** | **$280 \pm 10$** |
| **P94R/D160N/D165A/K190E/K190T**<br>**(saSrtA5M)** | **$0.04 \pm 0.02$** | **$47 \pm 11$** | **$770 \pm 120$** |
| **P94D/D160N/D165A/K190E/K190T**<br>**(P94D saSrtA5M)** | **$0.03 \pm 0.01$** | **$20 \pm 3$** | **$1500 \pm 200$** |

Our calculated kinetic parameters agreed with previously published values that the pentamutant mutations had a greater effect on *K*_m_ = 0.047 ± 0.011 mM (3.9-fold decrease as compared to wild-type) as compared to *k*_cat_ = 0.04 ± 0.02 s^-1^ (1.8-fold increase as compared to wild-type), although our results are not as dramatic as others reported.^44,61^ In general, we see a 6.4-fold increase in *k*_cat_/*K*_m_ for saSrtA5M (770 ± 120 M^-1^ s^-1^) as compared to wild-type saSrtA. Interestingly, while the *k*_cat_/*K*_m_ for P94R saSrtA is increased 2.3-fold as compared to wild-type (550 ± 30 M^-1^ s^-1^), the *k*_cat_/*K*_m_ for P94D saSrtA is increased 4.6-fold (**Table 1**). Consistent with this P94D *versus* P94R effect, we observed an additional 2-fold boost in *k*_cat_/*K*_m_ for P94D saSrtA5M (1500 ± 200 M^-1^ s^-1^) as compared to the saSrtA5M pentamutant (770 ± 120 M^-1^ s^-1^), which already contains the P94R mutation (**Table 1**). As noted above, these results were similar to our fluorescent peptide cleavage results (Figure 9); however, we were intrigued by the increased *k*_cat_/*K*_m_ for P94D saSrtA and P94D saSrtA5M versus P94R saSrtA and/or the pentamutant (with P94R) and wanted to further test these variants using additional HPLC-based assays.

Initially, we ran HPLC assays using hydroxylamine as our nucleophile; here, we are largely monitoring peptide cleavage analogous to our plate reader and enzyme kinetics assays. Indeed, we saw that the P94D saSrtA5M variant outperformed saSrtA5M for both the LP**E**TG and LP**K**TG substrates at all time points in our assay, with t=2 h substrate conversion values of 74.1 ± 3.8% (LP**E**TG) and 79.1 ± 3.3% (LP**K**TG) in triplicate experiments (Figures 11B-C**, S10A**). On average, these values were approximately 20-25% better than saSrtA5M with the LP**E**TG substrate (62.7 ± 1.7%, at t=2 h) (Figures 11B-C**, S10A**). Like our fluorescence assay results, SML efficiency was dramatically reduced for saSrtA5M with the LP**K**TG peptide, with 28.7 ± 0.01% substrate conversion at t=2 h (Figures 11B-C**, S10A**). Although saSrtA5M and P94D saSrtA5M outperformed the single P94D saSrtA mutation for the LP**E**TG substrate, product conversion was very similar at all time points for P94D saSrtA + LP**E**TG as compared to saSrtA5M + LP**K**TG. In our peptide cleavage assay, relative fluorescence units at t=2 h were similar for all P2 amino acids (A, E, K, S). Here, however, the P94D saSrtA enzyme showed a stronger preference for the LP**K**TG sequence (40.6 ± 1.5%, at t=2 h) over LP**E**TG, with a 1.4-fold increase in substrate conversion (Figures 6**, 11B-C, S10A**). This P2 preference for Lys by P94D saSrtA was also apparent when we normalized t=20 min RFU values to LP**E**TG in our peptide cleavage assay (Figure 5A). Overall, the SML experiments using peptide substrates revealed a greater increase in product formation with P94D saSrtA5M as compared to the P94R-containing saSrtA5M enzyme, although including the pentamutant background (i.e., additional four mutations, D160N/D165A/K190E/K196T) still provided the greatest cleavage efficiency difference, in the presence of mutation at P94, for the LP**E**TG peptide.

### Sortase mediated ligation reactions using a Gly-Gly nucleophile

Finally, we ran a sortase-mediated ligation reaction using a standard glycine (Gly-Gly) nucleophile. As described in the Materials and Methods, we used HPLC to measure reaction progress (as substrate conversion %) over 2 h with either an LP**E**TG or LP**K**TG substrate (full sequence: Abz-LP**X**TGGK(Dnp), as in other peptide assays) and an excess of diglycine (GG) peptide nucleophile. Here, we chose to compare the saSrtA5M and P94D saSrtA5M variants to see if the P94D mutation affected SML efficiency (**Figures 11D, S10A**). Consistent with our other results, while we saw a similar degree of cleavage product (GGK(Dnp)) formation at all time points for the LP**E**TG substrate, there was a substantial difference between saSrtA5M (46 ± 5%) as compared to P94D saSrtA5M (87 ± 2%) at t = 2 h using the LP**K**TG substrate (**Figures 11D, S10A**). In order to directly assess the formation of the desired ligation products (Abz-LP**X**TGG), the reactions were also characterized by LC-MS. In all cases clear signals for the ligation products were observed, with little to no hydrolysis product (<0.5% relative to ligation products) as estimated by mass spectrometry (representative mass spectrum shown in **Figure S10B**). These assays confirm that for certain LP**X**TG sequences (e.g., LP**K**TG) variants of saSrtA5M (e.g., P94D saSrtA5M) substantially outperform the original P94R-containing saSrtA5M enzyme in sortase-mediated ligation experiments. These experiments also demonstrate that the P94D mutation does not appear to have deleterious effects on the ratio of ligation product versus undesired substrate hydrolysis

## Discussion

Sortase-mediated ligation (SML) is a powerful technique used in a variety of protein engineering approaches.^26^ Historically, challenges in the utilization of SML included reversibility of the reaction, off-target hydrolysis of substrate, and a relatively low catalytic efficiency of the sortase catalytic domain.^26,57^ Within the SML engineering field, the one enzyme that is most widely used is saSrtA; specifically, engineered variants including a calcium-independent version of the pentamutant referenced here, termed the heptamutant, or saSrtA7M, as discussed above. We previously characterized a structurally-conserved loop in class A sortases, the β7-β8 loop, with a focus on *Streptococcus* SrtA enzymes.^36,51,52^ Analyses of experimental structures of saSrtA revealed an interaction between P94 and Y187 (in the β7-β8 loop) that is only present in the apo enzyme and which is not conserved in other SrtA enzymes (Figure 1C). We wanted to better understand the role of this position in saSrtA activity and specificity, and were additionally intrigued as P94 is a mutated residue in the catalytically-enhanced pentamutant saSrtA5M, which contains P94R.^44^

As previously observed, we predicted that mutation at P94 primarily affects the overall *K*_m_ of saSrtA. We found a wide range of relative activities for P94X variants and substrate peptides, with any mutation at this residue resulting in similar or better overall activity for all enzyme-substrate pairs tested (Figures 2**, 4**). In our data, the P94R variant did not emerge as one of the most active, and indeed, we hypothesize that the presence of the P2 Glu (LP**E**TG) may have influenced the original directed evolution results. The LP**E**TG sequence is derived from an endogenous target of saSrtA, the Spa protein, which may have evolved to optimize SrtA recognition and activity, as this was the most active sequence for almost all of our variants, presumably due to the R197 amino acid in saSrtA.

This Arg residue (R197 in saSrtA) is highly conserved in sortase enzymes. Traditionally, it is considered one of the catalytic triad residues. Recent work from ourselves and others suggested that while important, R197 likely does not stabilize the oxyanion intermediate in a catalytic manner, but instead contributes to substrate recognition and binding.^26,32–34^ This is consistent with the data presented here, wherein R197 stabilizes the substrate backbone in several AlphaFold3 models, as well as may influence P2 specificity. Indeed, the same group that developed saSrt5M later used the technique to engineer a sortase variant, SrtAβ, for use as a diagnostic and therapeutic tool for neurodegenerative disease.^27^ SrtAβ preferentially recognizes the LMVGG sequence and contains a Tyr at position 94 and Ser at the R197 position.^27^ The data presented here is consistent with these results, in that mutation of R197S presumably reduces the positive charge in the substrate binding cleft, potentially increasing affinity for a P2 Val. While P94Y saSrtA behaved like wild-type in our assays, the hydrophobic nature of the LMVGG sequence may be better tolerated.

Overall, our data argues that a combination of rational design based on structure-function and high throughput screening techniques may elucidate improved versions of SrtA for use in SML experiments. We were able to increase relative activity for certain substrates using pentamutant variants containing P94A and P94D mutations, and show that these variants (e.g., P94D saSrtA5M) can outperform saSrtA5M in SML experiments with certain LP**X**TG substrates (**Figure 11C,D**). Because previous work revealed that catalytic efficiency decreases in the heptamutant as compared to the pentamutant due to a 3-fold increase in *K*_m_, we did not test calcium-independent versions of our P94X variants here.^45,46^ We would expect a similar increase in relative *K*_m_ for any of the P94X variants tested; however, these effects provide additional support towards continued optimization of saSrtA variants for use in SML applications. Our data argues that a better understanding of the basic stereochemical mechanisms of SrtA activity and specificity can provide insight towards the design and development of next-generation tools in this wide-reaching field.

## Supporting information

Supplemental Information for Cox-Tigre et al.

## Acknowledgements

The authors would like to thank all members of the Amacher lab for scientific discussion and technical assistance. The authors would like to extend a special acknowledgement to Maria Jenkins for information technology (IT) support. This work was supported by NIH 1R15GM154315-01 to J.F. Amacher and J.M. Antos. It was additionally supported by NSF CHE-2044958 and a Cottrell Scholar Award from the Research Corporation for Science Advancement to J.F. Amacher, as well as a Beckman Scholars Award from the Arnold and Mabel Beckman Foundation to N. Cox-Tigre.

## References

(1) Rajagopal, M.; Walker, S. Envelope Structures of Gram-Positive Bacteria. Curr. Top. Microbiol. Immunol. 2017, 404, 1–44.

(2) Raz, A.; Fischetti, V. A. Sortase A Localizes to Distinct Foci on the *Streptococcus Pyogenes* Membrane. Proc. Natl. Acad. Sci. USA 2008, 105, 18549–18554.

(3) Ton-That, H.; Liu, G.; Mazmanian, S. K.; Faull, K. F.; Schneewind, O. Purification and Characterization of Sortase, the Transpeptidase That Cleaves Surface Proteins of *Staphylococcus Aureus* at the LPXTG Motif. Proc. Natl. Acad. Sci. USA 1999, 96, 12424–12429.

(4) Mazmanian, S. K.; Liu, G.; Ton-That, H.; Schneewind, O. *Staphylococcus Aureus* Sortase, an Enzyme That Anchors Surface Proteins to the Cell Wall. Science 1999, 285, 760–763.

(5) Ton-That, H.; Schneewind, O. Anchor Structure of *Staphylococcal* Surface Proteins. IV. Inhibitors of the Cell Wall Sorting Reaction. J. Biol. Chem. 1999, 274, 24316–24320.

(6) Zrelovs, N.; Kurbatska, V.; Rudevica, Z.; Leonchiks, A.; Fridmanis, D. Sorting out the Superbugs: Potential of Sortase A Inhibitors among Other Antimicrobial Strategies to Tackle the Problem of Antibiotic Resistance. Antibiotics (Basel) 2021, 10, 164.

(7) Jaudzems, K.; Kurbatska, V.; Je Kabsons, A.; Bobrovs, R.; Rudevica, Z.; Leonchiks, A. Targeting Bacterial Sortase A with Covalent Inhibitors: 27 New Starting Points for Structure-Based Hit-to-Lead Optimization. ACS Infect. Dis. 2020, 6, 186–194.

(8) Cascioferro, S.; Raffa, D.; Maggio, B.; Raimondi, M. V.; Schillaci, D.; Daidone, G. Sortase A Inhibitors: Recent Advances and Future Perspectives. J. Med. Chem. 2015, 58, 9108–9123.

(9) Thappeta, K. R. V.; Zhao, L. N.; Nge, C. E.; Crasta, S.; Leong, C. Y.; Ng, V.; Kanagasundaram, Y.; Fan, H.; Ng, S. B. In-Silico Identified New Natural Sortase A Inhibitors Disrupt S. Aureus Biofilm Formation. Int. J. Mol. Sci. 2020, 21(22):8601.

(10) Chan, A. H.; Wereszczynski, J.; Amer, B. R.; Yi, S. W.; Jung, M. E.; McCammon, J. A.; Clubb, R. T. Discovery of *Staphylococcus Aureus* Sortase A Inhibitors Using Virtual Screening and the Relaxed Complex Scheme. Chem Biol Drug Des 2013, 82, 418–428.

(11) Nitulescu, G.; Zanfirescu, A.; Olaru, O. T.; Nicorescu, I. M.; Nitulescu, G. M.; Margina, D. Structural Analysis of Sortase A Inhibitors. Molecules 2016, 21(11):1591.

(12) Maresso, A. W.; Wu, R.; Kern, J. W.; Zhang, R.; Janik, D.; Missiakas, D. M.; Duban, M.-E.; Joachimiak, A.; Schneewind, O. Activation of Inhibitors by Sortase Triggers Irreversible Modification of the Active Site. J. Biol. Chem. 2007, 282, 23129–23139.

(13) Wójcik, M.; Eleftheriadis, N.; Zwinderman, M. R. H.; Dömling, A. S. S.; Dekker, F. J.; Boersma, Y. L. Identification of Potential Antivirulence Agents by Substitution-Oriented Screening for Inhibitors of Streptococcus Pyogenes Sortase A. Eur. J. Med. Chem. 2019, 161, 93–100.

(14) Gosschalk, J. E.; Chang, C.; Sue, C. K.; Siegel, S. D.; Wu, C.; Kattke, M. D.; Yi, S. W.; Damoiseaux, R.; Jung, M. E.; Ton-That, H.;, et al. A Cell-Based Screen in *Actinomyces Oris* to Identify Sortase Inhibitors. Sci. Rep. 2020, 10, 8520.

(15) Zhang, J.; Liu, H.; Zhu, K.; Gong, S.; Dramsi, S.; Wang, Y.-T.; Li, J.; Chen, F.; Zhang, R.; Zhou, L.;, et al. Antiinfective Therapy with a Small Molecule Inhibitor of *Staphylococcus Aureus* Sortase. Proc. Natl. Acad. Sci. USA 2014, 111, 13517–13522.

(16) Volynets, G. P.; Barthels, F.; Hammerschmidt, S. J.; Moshynets, O. V.; Lukashov, S. S.; Starosyla, S. A.; Vyshniakova, H. V.; Iungin, O. S.; Bdzhola, V. G.; Prykhod’ko, A. O.;, et al. Identification of Novel Small-Molecular Inhibitors of *Staphylococcus Aureus* Sortase A Using Hybrid Virtual Screening. J Antibiot 2022, 75, 321–332.

(17) Sapra, R.; Rajora, A. K.; Kumar, P.; Maurya, G. P.; Pant, N.; Haridas, V. Chemical Biology of Sortase A Inhibition: A Gateway to Anti-Infective Therapeutic Agents. J. Med. Chem. 2021, 64, 13097–13130.

(18) Scott, C. J.; McDowell, A.; Martin, S. L.; Lynas, J. F.; Vandenbroeck, K.; Walker, B. Irreversible Inhibition of the Bacterial Cysteine Protease-Transpeptidase Sortase (SrtA) by Substrate-Derived Affinity Labels. Biochem. J. 2002, 366, 953–958.

(19) Kim, S.-W.; Chang, I.-M.; Oh, K.-B. Inhibition of the Bacterial Surface Protein Anchoring Transpeptidase Sortase by Medicinal Plants. Biosci. Biotechnol. Biochem. 2002, 66, 2751–2754.

(20) Obeng, E. M.; Fulcher, A. J.; Wagstaff, K. M. Harnessing Sortase A Transpeptidation for Advanced Targeted Therapeutics and Vaccine Engineering. Biotechnol. Adv. 2023, 64, 108108.

(21) Braga Emidio, N.; Cheloha, R. W. Semi-Synthetic Nanobody-Ligand Conjugates Exhibit Tunable Signaling Properties and Enhanced Transcriptional Outputs at Neurokinin Receptor-1. Protein Sci. 2024, 33, e4866.

(22) Zou, Z.; Ji, Y.; Schwaneberg, U. Empowering Site-Specific Bioconjugations In Vitro and In Vivo: Advances in Sortase Engineering and Sortase-Mediated Ligation. Angew. Chem. Int. Ed. 2023, e202310910.

(23) Beerli, R. R.; Hell, T.; Merkel, A. S.; Grawunder, U. Sortase Enzyme-Mediated Generation of Site-Specifically Conjugated Antibody Drug Conjugates with High In Vitro and In Vivo Potency. PLoS One 2015, 10, e0131177.

(24) Gébleux, R.; Briendl, M.; Grawunder, U.; Beerli, R. R. Sortase A Enzyme-Mediated Generation of Site-Specifically Conjugated Antibody-Drug Conjugates. Methods Mol. Biol. 2019, 2012, 1–13.

(25) Kumari, P.; Bowmik, S.; Paul, S. K.; Biswas, B.; Banerjee, S. K.; Murty, U. S.; Ravichandiran, V.; Mohan, U. Sortase A: A Chemoenzymatic Approach for the Labeling of Cell Surfaces. Biotechnol. Bioeng. 2021, 118, 4577–4589.

(26) Amacher, J. F.; Antos, J. M. Sortases: Structure, Mechanism, and Implications for Protein Engineering. Trends Biochem. Sci. 2024, 49, 596–610.

(27) Podracky, C. J.; An, C.; DeSousa, A.; Dorr, B. M.; Walsh, D. M.; Liu, D. R. Laboratory Evolution of a Sortase Enzyme That Modifies Amyloid-β Protein. Nat. Chem. Biol. 2021, 17, 317–325.

(28) Park, C.; Zhang, Y.; Jung, J. U.; Buron, L. D.; Lin, N.-P.; Hoeg-Jensen, T.; Chou, D. H.-C. Antagonistic Insulin Derivative Suppresses Insulin-Induced Hypoglycemia. J. Med. Chem. 2023, 66(11):7516–7522.

(29) Moliner-Morro, A.; J Sheward, D.; Karl, V.; Perez Vidakovics, L.; Murrell, B.; McInerney, G. M.; Hanke, L. Picomolar SARS-CoV-2 Neutralization Using Multi-Arm PEG Nanobody Constructs. Biomolecules 2020, 10(12):1661.

(30) Spirig, T.; Weiner, E. M.; Clubb, R. T. Sortase Enzymes in Gram-Positive Bacteria. Mol. Microbiol. 2011, 82, 1044–1059.

(31) Jacobitz, A. W.; Kattke, M. D.; Wereszczynski, J.; Clubb, R. T. Sortase Transpeptidases: Structural Biology and Catalytic Mechanism. Adv. Protein Chem. Struct. Biol. 2017, 109, 223–264.

(32) Tian, B.-X.; Eriksson, L. A. Catalytic Mechanism and Roles of Arg197 and Thr183 in the *Staphylococcus Aureus* Sortase A Enzyme. J. Phys. Chem. B 2011, 115, 13003–13011.

(33) Johnson, D. A.; Piper, I. M.; Vogel, B. A.; Jackson, S. N.; Svendsen, J. E.; Kodama, H. M.; Lee, D. E.; Lindblom, K. M.; McCarty, J.; Antos, J. M.;, et al. Structures of *Streptococcus Pyogenes* Class A Sortase in Complex with Substrate and Product Mimics Provide Key Details of Target Recognition. J. Biol. Chem. 2022, 298, 102446.

(34) Chen, J.-L.; Wang, X.; Yang, F.; Li, B.; Otting, G.; Su, X.-C. 3D Structure of the Transient Intermediate of the Enzyme–substrate Complex of Sortase A Reveals How Calcium Binding and Substrate Recognition Cooperate in Substrate Activation. ACS Catal. 2023, 13, 11610–11624.

(35) Malik, A.; Kim, S. B. A Comprehensive in Silico Analysis of Sortase Superfamily. J. Microbiol. 2019, 57, 431–443.

(36) Valgardson, J. D.; Struyvenberg, S. A.; Sailer, Z. R.; Piper, I. M.; Svendsen, J. E.; Johnson, D. A.; Vogel, B. A.; Antos, J. M.; Harms, M. J.; Amacher, J. F. Comparative Analysis and Ancestral Sequence Reconstruction of Bacterial Sortase Family Proteins Generates Functional Ancestral Mutants with Different Sequence Specificities. Bacteria 2022, 1, 121–135.

(37) Nikghalb, K. D.; Horvath, N. M.; Prelesnik, J. L.; Banks, O. G. B.; Filipov, P. A.; Row, R. D.; Roark, T. J.; Antos, J. M. Expanding the Scope of Sortase-Mediated Ligations by Using Sortase Homologues. Chembiochem 2018, 19, 185–195.

(38) Schmohl, L.; Bierlmeier, J.; von Kügelgen, N.; Kurz, L.; Reis, P.; Barthels, F.; Mach, P.; Schutkowski, M.; Freund, C.; Schwarzer, D. Identification of Sortase Substrates by Specificity Profiling. Bioorg. Med. Chem. 2017, 25, 5002–5007.

(39) Ilangovan, U.; Ton-That, H.; Iwahara, J.; Schneewind, O.; Clubb, R. T. Structure of Sortase, the Transpeptidase That Anchors Proteins to the Cell Wall of *Staphylococcus Aureus*. Proc. Natl. Acad. Sci. USA 2001, 98, 6056–6061.

(40) Suree, N.; Liew, C. K.; Villareal, V. A.; Thieu, W.; Fadeev, E. A.; Clemens, J. J.; Jung, M. E.; Clubb, R. T. The Structure of the *Staphylococcus Aureus* Sortase-Substrate Complex Reveals How the Universally Conserved LPXTG Sorting Signal Is Recognized. J. Biol. Chem. 2009, 284, 24465–24477.

(41) Chan, A. H.; Yi, S. W.; Terwilliger, A. L.; Maresso, A. W.; Jung, M. E.; Clubb, R. T. Structure of the *Bacillus Anthracis* Sortase A Enzyme Bound to Its Sorting Signal: A FLEXIBLE AMINO-TERMINAL APPENDAGE MODULATES SUBSTRATE ACCESS. J. Biol. Chem. 2015, 290, 25461–25474.

(42) Weiner, E. M.; Robson, S.; Marohn, M.; Clubb, R. T. The Sortase A Enzyme That Attaches Proteins to the Cell Wall of *Bacillus Anthracis* Contains an Unusual Active Site Architecture. J. Biol. Chem. 2010, 285, 23433–23443.

(43) Race, P. R.; Bentley, M. L.; Melvin, J. A.; Crow, A.; Hughes, R. K.; Smith, W. D.; Sessions, R. B.; Kehoe, M. A.; McCafferty, D. G.; Banfield, M. J. Crystal Structure of *Streptococcus Pyogenes* Sortase A: Implications for Sortase Mechanism. J. Biol. Chem. 2009, 284, 6924–6933.

(44) Chen, I.; Dorr, B. M.; Liu, D. R. A General Strategy for the Evolution of Bond-Forming Enzymes Using Yeast Display. Proc. Natl. Acad. Sci. USA 2011, 108, 11399–11404.

(45) Hirakawa, H.; Ishikawa, S.; Nagamune, T. Design of Ca2+-Independent *Staphylococcus Aureus* Sortase A Mutants. Biotechnol. Bioeng. 2012, 109, 2955–2961.

(46) Hirakawa, H.; Ishikawa, S.; Nagamune, T. Ca2+ - Independent Sortase-A Exhibits High Selective Protein Ligation Activity in the Cytoplasm of *Escherichia Coli*. Biotechnol. J. 2015, 10, 1487–1492.

(47) Witte, M. D.; Wu, T.; Guimaraes, C. P.; Theile, C. S.; Blom, A. E. M.; Ingram, J. R.; Li, Z.; Kundrat, L.; Goldberg, S. D.; Ploegh, H. L. Site-Specific Protein Modification Using Immobilized Sortase in Batch and Continuous-Flow Systems. Nat. Protoc. 2015, 10, 508–516.

(48) Bentley, M. L.; Gaweska, H.; Kielec, J. M.; McCafferty, D. G. Engineering the Substrate Specificity of Staphylococcus Aureus Sortase A. The Beta6/Beta7 Loop from SrtB Confers NPQTN Recognition to SrtA. J. Biol. Chem. 2007, 282, 6571–6581.

(49) Ugur, I.; Schatte, M.; Marion, A.; Glaser, M.; Boenitz-Dulat, M.; Antes, I. Ca2+ Binding Induced Sequential Allosteric Activation of Sortase A: An Example for Ion-Triggered Conformational Selection. PLoS One 2018, 13, e0205057.

(50) Piotukh, K.; Geltinger, B.; Heinrich, N.; Gerth, F.; Beyermann, M.; Freund, C.; Schwarzer, D. Directed Evolution of Sortase A Mutants with Altered Substrate Selectivity Profiles. J. Am. Chem. Soc. 2011, 133, 17536–17539.

(51) Piper, I. M.; Struyvenberg, S. A.; Valgardson, J. D.; Johnson, D. A.; Gao, M.; Johnston, K.; Svendsen, J. E.; Kodama, H. M.; Hvorecny, K. L.; Antos, J. M.;, et al. Sequence Variation in the Β7-Β8 Loop of Bacterial Class A Sortase Enzymes Alters Substrate Selectivity. J. Biol. Chem. 2021, 297, 100981.

(52) Gao, M.; Johnson, D. A.; Piper, I. M.; Kodama, H. M.; Svendsen, J. E.; Tahti, E.; Longshore-Neate, F.; Vogel, B.; Antos, J. M.; Amacher, J. F. Structural and Biochemical Analyses of Selectivity Determinants in Chimeric *Streptococcus* Class A Sortase Enzymes. Protein Sci. 2022, 31, 701–715.

(53) Kodama, H. M.; Lindblom, K. M.; Walkenhauer, E. G.; Antos, J. M.; Amacher, J. F. Amino Acid Variability at W194 of *Staphylococcus Aureus* Sortase A Alters Nucleophile Specificity. Protein Sci. 2024, 33, e5212.

(54) Abramson, J.; Adler, J.; Dunger, J.; Evans, R.; Green, T.; Pritzel, A.; Ronneberger, O.; Willmore, L.; Ballard, A. J.; Bambrick, J.;, et al. Accurate Structure Prediction of Biomolecular Interactions with AlphaFold 3. Nature 2024, 630, 493–500.

(55) Gao, X.; Dong, X.; Li, X.; Liu, Z.; Liu, H. Prediction of Disulfide Bond Engineering Sites Using a Machine Learning Method. Sci. Rep. 2020, 10, 10330.

(56) Vogel, B. A.; Blount, J. M.; Kodama, H. M.; Goodwin-Rice, N. J.; Andaluz, D. J.; Jackson, S. N.; Antos, J. M.; Amacher, J. F. A Unique Binding Mode of P1’ Leu-Containing Target Sequences for *Streptococcus Pyogenes* Sortase A Results in Alternative Cleavage. *RSC Chem*. Biol. 2024, 5, 30–40.

(57) Morgan, H. E.; Turnbull, W. B.; Webb, M. E. Challenges in the Use of Sortase and Other Peptide Ligases for Site-Specific Protein Modification. Chem. Soc. Rev. 2022, 51, 4121–4145.

(58) Frankel, B. A.; Kruger, R. G.; Robinson, D. E.; Kelleher, N. L.; McCafferty, D. G. *Staphylococcus Aureus* Sortase Transpeptidase SrtA: Insight into the Kinetic Mechanism and Evidence for a Reverse Protonation Catalytic Mechanism. Biochemistry 2005, 44, 11188–11200.

(59) Huang, X.; Aulabaugh, A.; Ding, W.; Kapoor, B.; Alksne, L.; Tabei, K.; Ellestad, G. Kinetic Mechanism of *Staphylococcus Aureus* Sortase SrtA. Biochemistry 2003, 42, 11307–11315.

(60) Kruger, R. G.; Otvos, B.; Frankel, B. A.; Bentley, M.; Dostal, P.; McCafferty, D. G. Analysis of the Substrate Specificity of the *Staphylococcus Aureus* Sortase Transpeptidase SrtA. Biochemistry 2004, 43, 1541–1551.

(61) Wang, C.; Desmet, R.; Snella, B.; Vicogne, J.; Melnyk, O.; Agouridas, V. Leveraging Sortase A Electrostatics for Powerful Transpeptidation Reactions. Angew. Chem. Int. Ed. 2025, 64, e202507236.

(62) Chen, L.; Cohen, J.; Song, X.; Zhao, A.; Ye, Z.; Feulner, C. J.; Doonan, P.; Somers, W.; Lin, L.; Chen, P. R. Improved Variants of SrtA for Site-Specific Conjugation on Antibodies and Proteins with High Efficiency. Sci. Rep. 2016, 6, 31899.

(63) Bentley, M. L.; Lamb, E. C.; McCafferty, D. G. Mutagenesis Studies of Substrate Recognition and Catalysis in the Sortase A Transpeptidase from *Staphylococcus Aureus*. J. Biol. Chem. 2008, 283, 14762–14771.

