## Supplemental Information for Cox-Tigre et al. for "Amino acid variants at the P94 position in *Staphylococcus aureus* class A sortase modulate substrate binding and enzyme activity"

**Table of Contents**

**Supplemental Results: AlphaFold3 modeling.** 2

**Table S1. Mass spectrometry analyses of saSrtA variants and synthesized peptides.** 3

**Table S2. AlphaFold3 output data (pTM and ipTM, if relevant).** 4

**Table S3. β7-β8 loop flexibility in AlphaFold3 models, ± $Ca^{2+}$.** 5

**Table S4. Initial velocities (in RFU/min) for all enzymes/substrates tested.** 6

**Figure S1. SDS-PAGE gel for purified saSrtA variants.** 7

**Figure S2. Fluorescence data for saSrtA variants with LPXTG substrates (t=2 h).** 8

**Figure S3. AlphaFold3 models of P94X saSrtA variants with LPETG substrate bound, including wild-type saSrtA and distance measurements.** 9

**Figure S4. AlphaFold3 model of saSrtA-LPETG with peptide outside binding pocket.** 10

**Figure S5. AlphaFold3 models of P94X saSrtA variants, ± $Ca^{2+}$.** 11

**Figure S6. AlphaFold3 output data.** 13

**Figure S7. Initial velocity calculations for P94X saSrtA variants.** 17

**Figure S8. Comparison of initial velocities for P94X saSrtA variants.** 18

**Figure S9. Replicate data for enzyme kinetics assays.** 19

**Figure S10. Representative HPLC traces for assays with saSrtA5M, P94D saSrtA5M, and P94D saSrtA ($H_2NOH$ and/or Gly-Gly nucleophiles).** 20

*AlphaFold3 modeling of P94X saSrtA variants.* We used AlphaFold3 to model the WT and P94R saSrtA enzymes with LPETGG and $Ca^{2+}$, as well as the P94G, P94S, and P94T variants which all showed >3-fold increases in relative activity as compared to WT (**Figures S3A, Table S2**).[52] Interestingly, for all the saSrtA variants modeled, only the WT enzyme was found to not properly bind the substrate in our initial models, with the β7-β8 loop adopting the 'closed, down' conformation (**Figure S4**). We reasoned this may reflect the hypothesized increase in relative $K_m$ as compared to the P94 mutants. Therefore, we added an additional three N-terminal residues, derived from the endogenous substrate Spa (UniProt ID SPA_STAAE), AQALPETGG, for our subsequent modeling, which did allow for successful substrate binding to WT saSrtA (**Figure S3A**). In general, there are no interactions between side chain atoms in the WT or P94X variants and the peptide which would explain the differences in activity we observed.

We next used AlphaFold3 to model all P94X variants alone in both the -$Ca^{2+}$ and +$Ca^{2+}$ states, in order to better characterize the non-substrate bound enzymes (**Table S2**). Our -$Ca^{2+}$ models were largely consistent with the loop 'closed, down' conformation, with average distances between the $C_\alpha$ atoms of P94X and Y187 for the 5 output AlphaFold3 models ranging from 7.0 ± 0.2 Å (P94S) to 10.8 ± 3.9 Å (P94Y) (**Figures S3, S5-S6, and Table S3**). While there was higher variability in the +$Ca^{2+}$ models, which should represent an allosterically activated saSrtA conformation, measured distances still trended towards a more closed conformation. Here, while the WT output models maintained the loop 'down' conformation (7.6 ± 0.1 Å), larger average distances were observed for the P94X variants, with P94D (11.5 ± 2.3 Å) and P94Y (11.6 ± 4.2 Å) showing the largest differences (**Figures S3, S5-S6, Table S3**). Overall, there were no observed differences in this distance between P94X variants, despite our original prediction that these models could be used to infer relative flexibility differences upon P94X mutation. However, as these analyses are based on structural models, it would be interesting to investigate relative loop flexibility experimentally and/or to use molecular dynamics simulations to probe the AlphaFold3 models on a deeper level.

**Table S1. Mass spectrometry analysis of all enzyme variants and peptide substrates / peptide reaction products.** Predicted masses for enzyme variants were calculated using ExPasy ProtParam. Predicted and observed masses for protein variants represent average molecular weight (MW). Predicted and observed masses for peptides represent $[M+H]^+$ ions (monoisotopic). All peptide substrates (Abz-LP**X**TGGK(Dnp)) were synthesized with a C-terminal primary amide ($-NH_2$). Peptide ligation products (Abz-LP**X**TGG) contain a C-terminal carboxylic acid (-COOH).

| P94X | Predicted Mass (Da) | Observed Mass (Da) |
|---|---|---|
| A | 18710.0 | 18711.7 |
| D | 18754.0 | 18756.2 |
| E | 18768.1 | 18770.7 |
| F | 18786.1 | 18789.5 |
| G | 18696.0 | 18699.6 |
| H | 18776.1 | 18779.5 |
| I | 18752.1 | 18756.3 |
| K | 18767.1 | 18768.8 |
| L | 18752.1 | 18757.9 |
| M | 18770.1 | 18773.7 |
| N | 18753.1 | 18757.8 |
| WT (P) | 18736.1 | 18738.7 |
| Q | 18767.1 | 18771.4 |
| R | 18795.1 | 18799.6 |
| S | 18726.0 | 18730.3 |
| T | 18740.1 | 18739.7 |
| V | 18738.1 | 18741.4 |
| W | 18825.2 | 18829.1 |
| Y | 18802.1 | 18805.3 |
| saSrtA5M | 18724.0 | 18721.7 |
| P94A 5M | 18638.9 | 18636.7 |
| P94D 5M | 18682.9 | 18681.9 |
| **Peptides** | | |
| Abz-LP**A**TGGK(Dnp) | 927.4 | 927.5 |
| Abz-LP**E**TGGK(Dnp) | 985.4 | 985.3 |
| Abz-LP**K**TGGK(Dnp) | 984.5 | 984.6 |
| Abz-LP**S**TGGK(Dnp) | 943.4 | 943.6 |
| GGK(Dnp) | 426.2 | 426.1 |
| Abz-LP**E**TGG | 692.3 | 692.3 |
| Abz-LP**K**TGG | 691.4 | 691.4 |

**Table S2. AlphaFold3 output data (pTM and ipTM, if relevant).**

| Variant | Calcium (X=yes) | Ligand | pTM | ipTM |
|---|---|---|---|---|
| WT (P94) | | | | |
| WT (P94) | X | | 0.94 | 0.88 |
| WT (P94) | X | LPETGG | 0.87 | 0.62 |
| WT (P94) | X | AQALPETGG | 0.88 | 0.57 |
| P94A | | | 0.84 | |
| P94A | X | | 0.85 | 0.95 |
| P94D | | | 0.85 | |
| P94D | X | | 0.84 | 0.95 |
| P94D | X | AQALPATGG | 0.89 | 0.58 |
| P94E | | | 0.85 | |
| P94E | X | | 0.85 | 0.95 |
| P94E | X | AQALPATGG | 0.89 | 0.59 |
| P94F | | | 0.83 | |
| P94F | X | | 0.85 | 0.95 |
| P94G | | | 0.84 | |
| P94G | X | | 0.85 | 0.95 |
| P94H | | | 0.83 | |
| P94H | X | | 0.85 | 0.95 |
| P94I | | | 0.84 | |
| P94I | X | | 0.86 | 0.95 |
| P94K | | | 0.84 | |
| P94K | X | | 0.86 | 0.95 |
| P94L | | | 0.83 | |
| P94L | X | | 0.85 | 0.95 |
| P94M | | | 0.84 | |
| P94M | X | | 0.85 | 0.95 |
| P94N | | | 0.84 | |
| P94N | X | | 0.85 | 0.94 |
| P94Q | | | 0.85 | |
| P94Q | X | | 0.86 | 0.95 |
| P94R | | | 0.85 | |
| P94R | X | | 0.86 | 0.95 |
| P94R | X | AQALPATGG | 0.88 | 0.59 |
| P94S | | | 0.85 | |
| P94S | X | | 0.8§ | 0.95 |
| P94T | | | 0.85 | |
| P94T | X | | 0.85 | 0.95 |
| P94V | | | 0.84 | |
| P94V | X | | 0.85 | 0.95 |
| P94W | | | 0.83 | |
| P94W | X | | 0.85 | 0.95 |
| P94Y | | | 0.82 | |
| P94Y | X | | 0.85 | 0.95 |
| P94G | X | LPETGG | 0.88 | 0.61 |
| P94R | X | LPETGG | 0.85 | 0.56 |
| P94S | X | LPETGG | 0.86 | 0.64 |
| P94T | X | LPETGG | 0.87 | 0.6 |

**Table S3. β7-β8 loop flexibility in AlphaFold3 models, ± $Ca^{2+}$.** Relative flexibility was suggested by measuring the distance (in Å) between the Cα atoms P94X and Y187 in 5 output AlphaFold3 models, then averaging these values. The wild-type (WT) saSrtA models were used as benchmarks for relatively “little to no” flexibility, as these distances closely match those of the experimental structure, PDB ID 1IJA.

### -$Ca^{2+}$ models, distances measured using PyMOL (in Å)

| | P94A | P94D | P94E | P94F | P94G | P94H | P94I | P94K | P94L | P94M | P94N | WT | P94Q | P94R | P94S | P94T | P94V | P94W | P94Y |
|---|---|---|---|---|---|---|---|---|---|---|---|---|---|---|---|---|---|---|---|
| AF3 model_0 | 6.8 | 6.9 | 7.1 | 7.2 | 7.1 | 7.1 | 7.3 | 7.5 | 7.4 | 7.2 | 7.9 | 7.4 | 7.2 | 7.1 | 6.8 | 7.3 | 7.5 | 7.2 | 8.5 |
| AF3 model_1 | 6.8 | 6.9 | 7.3 | 7.4 | 7.1 | 8.2 | 7.3 | 7.4 | 7.7 | 7.2 | 6.8 | 7.5 | 7.4 | 7.7 | 7 | 7.3 | 7.3 | 7.1 | 7.5 |
| AF3 model_2 | 7 | 7 | 7.1 | 7.2 | 6.9 | 9.4 | 7.7 | 7.1 | 8.4 | 7.3 | 7 | 7.4 | 7.2 | 7.4 | 6.9 | 7.6 | 8.2 | 8 | 9.3 |
| AF3 model_3 | 7 | 7.1 | 7.1 | 8.6 | 7.3 | 9.8 | 7.9 | 7.3 | 10.1 | 8 | 8.4 | 7.5 | 7.5 | 7.4 | 7.2 | 7.2 | 8.7 | 8.3 | 11.3 |
| AF3 model_4 | 7.8 | 7.7 | 7.8 | 10.1 | 9.2 | 12.2 | 8.3 | 9.2 | 8.4 | 9.3 | 8 | 8.1 | 7.4 | 7.2 | 6.9 | 7.2 | 9.9 | 8.3 | 17.3 |
| **Average** | **7.08** | **7.12** | **7.28** | **8.1** | **7.52** | **9.34** | **7.7** | **7.7** | **8.4** | **7.8** | **7.62** | **7.58** | **7.34** | **7.36** | **6.96** | **7.32** | **8.32** | **7.78** | **10.78** |
| **Std. Dev.** | **0.4147288** | **0.334664** | **0.303315** | **1.260952** | **0.9497368** | **1.9178113** | **0.4242641** | **0.8514693** | **1.0464225** | **0.9027735** | **0.6870226** | **0.2949576** | **0.1341641** | **0.2302173** | **0.1516575** | **0.1643168** | **1.044988** | **0.5890671** | **3.9028195** |

### +$Ca^{2+}$ models, distances measured using PyMOL (in Å)

| | P94A_Ca | P94D_Ca | P94E_Ca | P94F_Ca | P94G_Ca | P94H_Ca | P94I_Ca | P94K_Ca | P94L_Ca | P94M_Ca | P94N_Ca | WT_Ca | P94Q_Ca | P94R_Ca | P94S_Ca | P94T_Ca | P94V_Ca | P94W_Ca | P94Y_Ca |
|---|---|---|---|---|---|---|---|---|---|---|---|---|---|---|---|---|---|---|---|
| AF3 model_0 | 12.2 | 12 | 7.8 | 10.7 | 10.9 | 10 | 7.7 | 7.9 | 8.7 | 8.6 | 7.3 | 7.5 | 7.6 | 8.2 | 6.6 | 8 | 8.5 | 7.3 | 8.3 |
| AF3 model_1 | 8.8 | 7.8 | 8 | 10.9 | 10.2 | 12.8 | 8.4 | 8 | 8 | 7.1 | 7.8 | 7.5 | 9.3 | 11.8 | 7.3 | 8.9 | 8.6 | 8 | 17.6 |
| AF3 model_2 | 8.3 | 12.4 | 7.4 | 12.6 | 8.5 | 9.4 | 8.8 | 8.5 | 8.1 | 7.9 | 9.5 | 7.6 | 8.7 | 13.3 | 8 | 7.9 | 8 | 8 | 10 |
| AF3 model_3 | 8.5 | 11.1 | 8.2 | 12.4 | 11.4 | 10.8 | 7.7 | 8.1 | 7.5 | 7.8 | 7.4 | 7.6 | 7.8 | 12.1 | 7.1 | 14.5 | 8.8 | 10.7 | 7.9 |
| AF3 model_4 | 9.9 | 14.1 | 10.3 | 8 | 11.6 | 8.4 | 8.9 | 13.1 | 7.7 | 8.4 | 14.8 | 7.7 | 7.9 | 9.3 | 9.5 | 7.7 | 13.9 | 7.3 | 14.1 |
| **Average** | **9.54** | **11.48** | **8.34** | **10.92** | **10.52** | **10.28** | **8.3** | **9.12** | **8** | **7.96** | **9.36** | **7.58** | **8.26** | **10.94** | **7.7** | **9.4** | **9.56** | **8.26** | **11.58** |
| **Std. Dev.** | **1.6102795** | **2.327445** | **1.1349009** | **1.8430952** | **1.2517987** | **1.6589153** | **0.5787918** | **2.2365151** | **0.4582576** | **0.585662** | **3.1674911** | **0.083666** | **0.7162402** | **2.1125814** | **1.1247222** | **2.8879058** | **2.4439722** | **1.4081903** | **4.164973** |

**Table S4. Initial velocities (in RFU/min) for all enzymes/substrates tested.** Data used for calculations is shown in **Figure S7**. Linear regression curves were calculated using GraphPad Prism. All $R^2$ values were $\geq 0.996$, with most $\geq 0.999$.

| Substrate | LPATG | | LPETG | | LPKTG | | LPSTG | |
|---|---|---|---|---|---|---|---|---|
| | Initial Velocity | Standard Deviation | Initial Velocity | Standard Deviation | Initial Velocity | Standard Deviation | Initial Velocity | Standard Deviation |
| **P94A** | 2834.3 | 77.1 | 3760.0 | 65.6 | 1788.0 | 159.8 | 2749.0 | 109.3 |
| **P94D** | 3671.0 | 208.4 | 2968.3 | 105.7 | 3609.7 | 40.0 | 3803.0 | 110.8 |
| **P94E** | 2943.7 | 6.1 | 2469.3 | 161.6 | 3107.3 | 152.6 | 2930.3 | 116.6 |
| **P94F** | 786.2 | 20.3 | 871.7 | 63.6 | 480.5 | 34.3 | 662.5 | 26.0 |
| **P94G** | 3284.7 | 88.3 | 4310.0 | 206.9 | 2405.0 | 82.6 | 2972.3 | 229.6 |
| **P94H** | 1922.0 | 123.8 | 4012.7 | 173.3 | 1334.7 | 21.6 | 2172.0 | 60.2 |
| **P94I** | 1359.7 | 21.4 | 1592.3 | 58.0 | 877.5 | 17.4 | 1139.0 | 34.6 |
| **P94K** | 1168.7 | 34.8 | 2752.3 | 142.3 | 606.0 | 50.4 | 1090.0 | 20.7 |
| **P94L** | 1070.5 | 98.9 | 1546.3 | 16.3 | 783.0 | 45.9 | 1140.7 | 71.8 |
| **P94M** | 1098.7 | 50.8 | 1364.3 | 27.2 | 845.2 | 50.8 | 1036.3 | 24.6 |
| **P94N** | 1826.0 | 47.7 | 2865.0 | 61.3 | 1349.3 | 5.1 | 2172.0 | 77.7 |
| **WT (P94)** | 598.2 | 24.8 | 937.8 | 47.6 | 467.2 | 14.8 | 610.0 | 6.3 |
| **P94Q** | 1726.0 | 61.0 | 2971.3 | 250.8 | 1397.7 | 15.3 | 1717.7 | 37.6 |
| **P94R** | 722. 7 | 25.5 | 2140.0 | 49.8 | 405.2 | 34.9 | 901.3 | 43.8 |
| **P94S** | 2764.3 | 27.6 | 4214.7 | 133.1 | 2326.7 | 54.5 | 3158.3 | 148.0 |
| **P94T** | 3428.3 | 37.6 | 4788.0 | 125.0 | 2909.0 | 64.1 | 3568.7 | 95.3 |
| **P94V** | 1172.0 | 56.3 | 1636.3 | 97.1 | 950.3 | 34.6 | 1187.2 | 179.3 |
| **P94W** | 875.2 | 31.2 | 1400.3 | 68.5 | 624.5 | 30.8 | 716.3 | 29.8 |
| **P94Y** | 987.3 | 48.3 | 1320.0 | 60.1 | 630.7 | 0.8 | 779.8 | 18.2 |
| **saSrtA5M** | 520.7 | 50.9 | 1285.3 | 58.6 | 339.7 | 15.0 | 617.5 | 21.8 |
| **P94A 5M** | 730.8 | 24.8 | 1472.0 | 34.9 | 711.7 | 18.7 | 859.2 | 59.5 |
| **P94D 5M** | 846.3 | 12.3 | 1128.0 | 52.0 | 1107.0 | 22.3 | 931.2 | 99.3 |

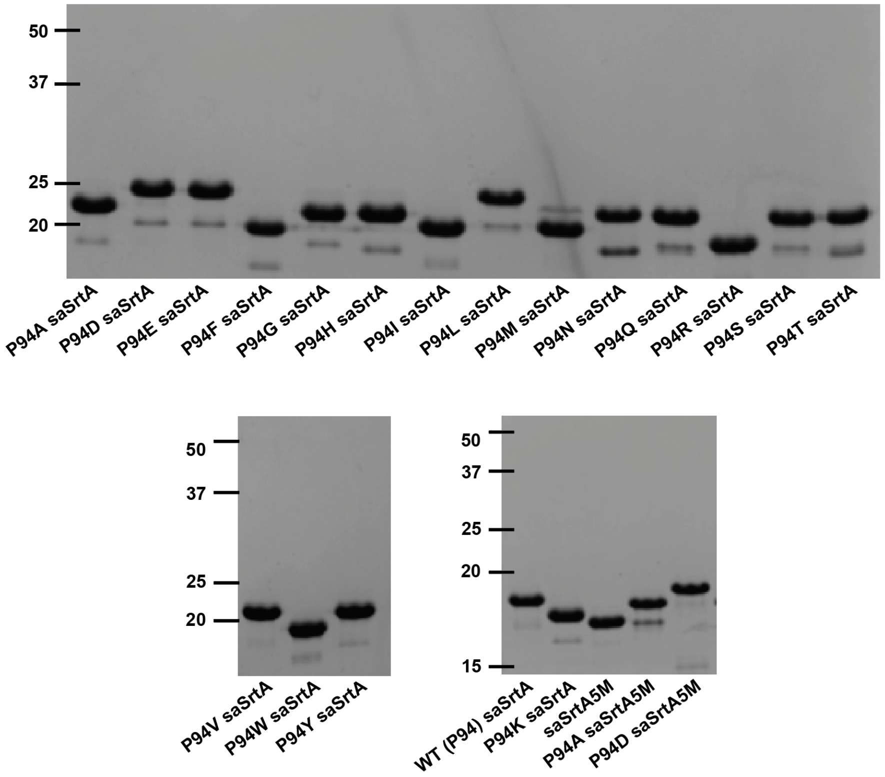

**Figure S1. SDS-PAGE gel for purified saSrtA variants.** SDS-PAGE gel of P94X saSrtA variants, following size exclusion chromatography (SEC), to show relative purity of enzymes used.

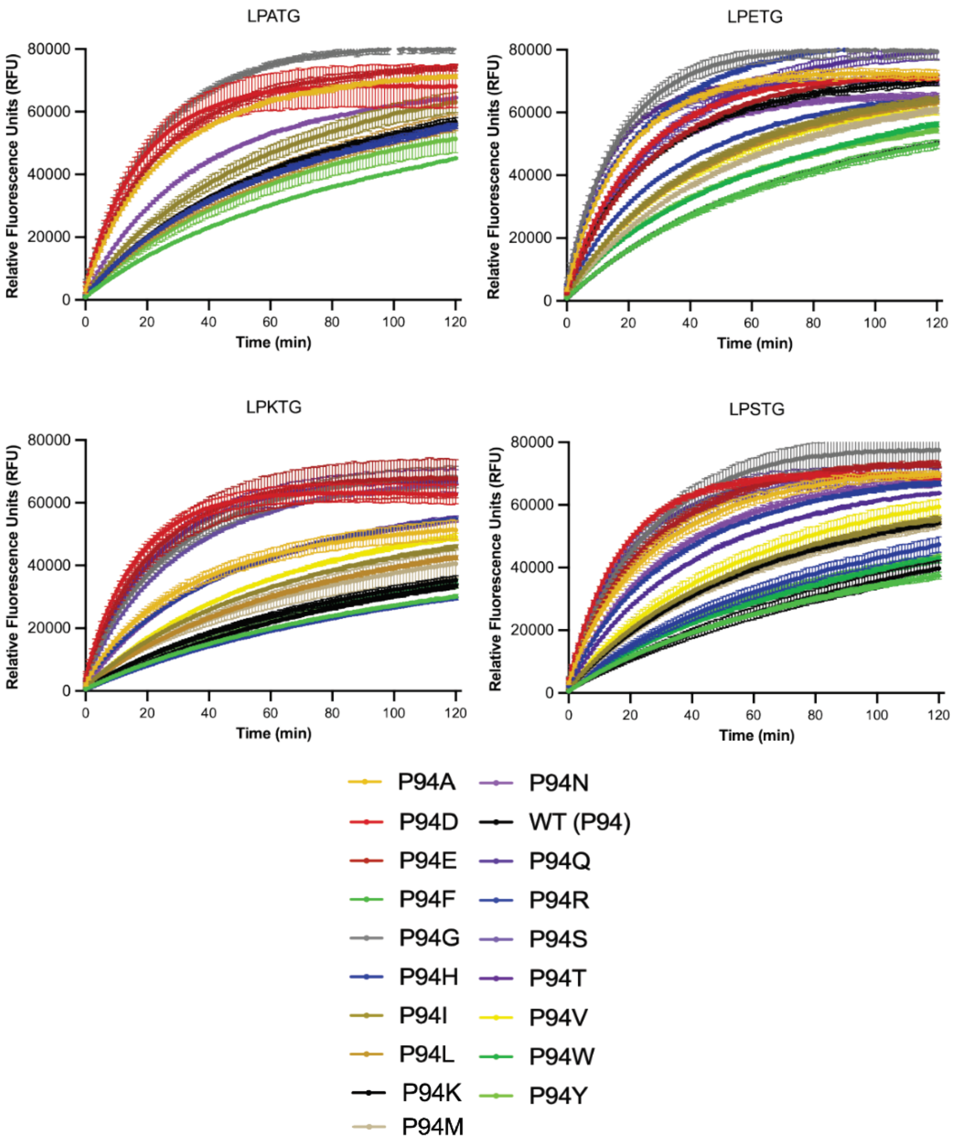

**Figure S2. Fluorescence data for saSrtA variants with LPXTG substrates (t=2 h).** The entire time course (t=2 h) for the activity assays is shown. This data matches that shown for t=60 min in **Figures 2A and 4**. Data is shown as the average of triplicate assays, with error bars equal to the standard deviation for each time point.

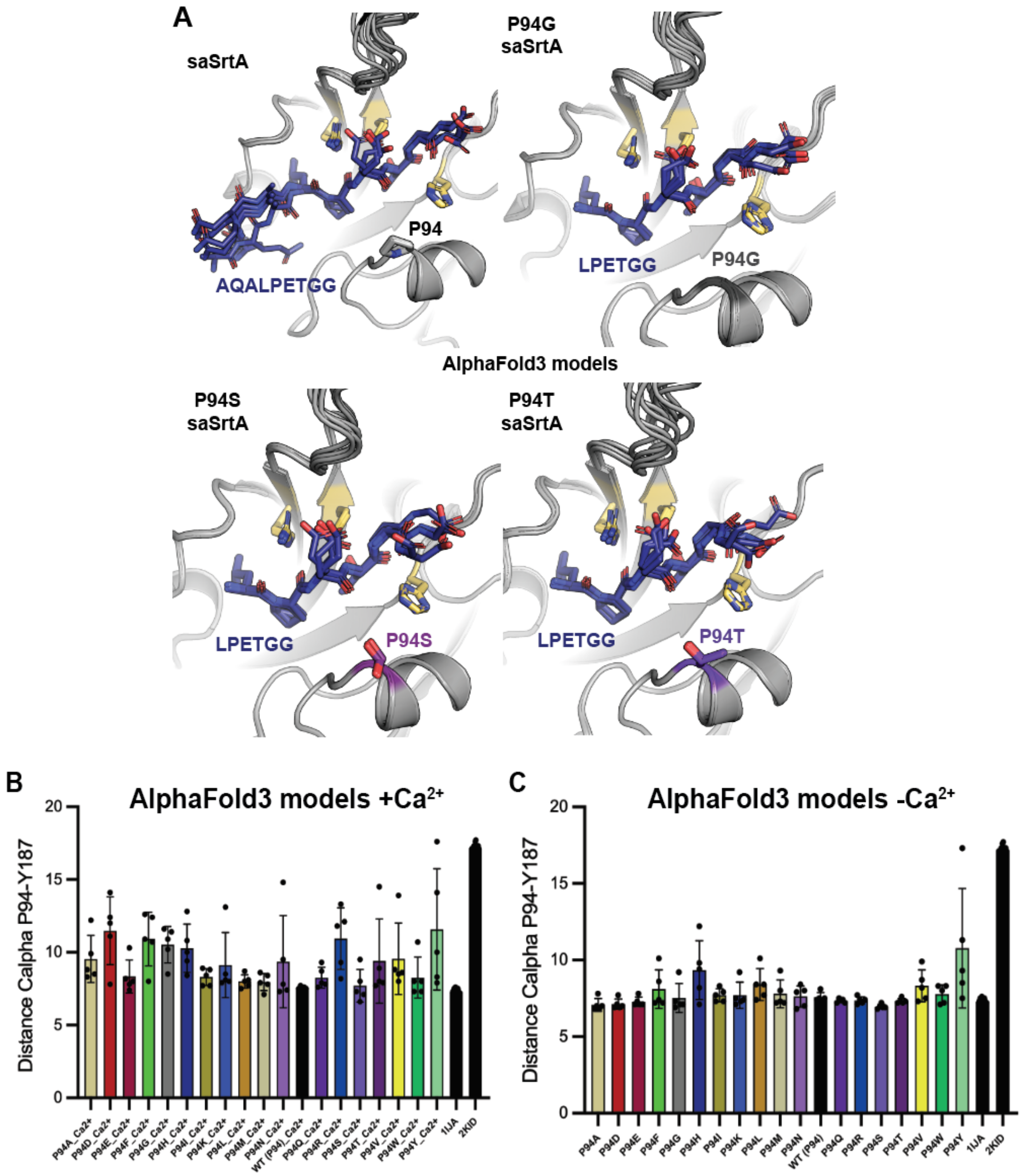

**Figure S3. AlphaFold3 models of P94X saSrtA variants with LPETG substrate bound, including wild-type saSrtA and distance measurements.** (**A**) AlphaFold3 models of saSrtA-AQALPETGG, P94G saSrtA-LPETGG, P94S saSrtA-LPETGG, and P94T saSrtA-LPETGG, as labeled. For all, the enzyme is in gray cartoon, with catalytic residue side chains as yellow sticks, and the P94X position as side chain sticks and colored as in **Figure 2**. The peptides are shown as blue sticks and colored by heteroatom. For all, C=as described, O=red, N=blue, S=gold. (**B-C**) Bar graph measurements, displaying average values ± standard deviation for the distance between P94X-Y187 C$\alpha$ atoms for all AlphaFold3 models of P94X saSrtA variants +$Ca^{2+}$ (**B**) or -$Ca^{2+}$ (**C**), as labeled. Measurements from experimental structures are also shown (right-most black bars). There were no observed differences in distances between P94X variants for either the +$Ca^{2+}$ or the -$Ca^{2+}$ models. AlphaFold3 output data is in **Tables S2-3** and **Figures S5-S6.**

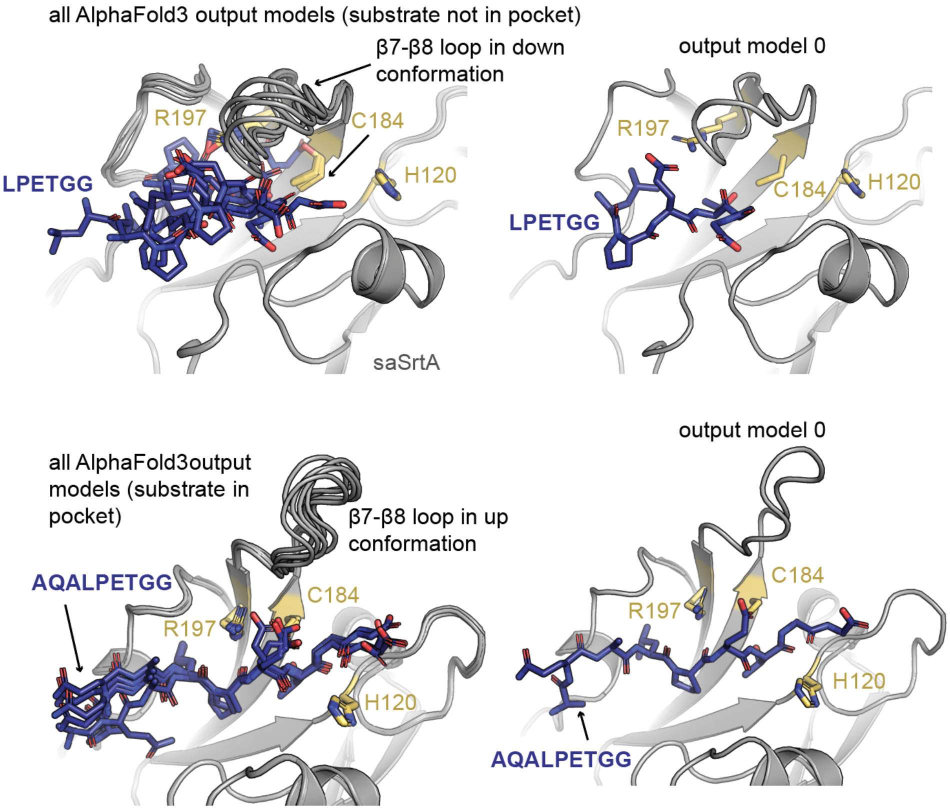

**Figure S4. AlphaFold3 model of saSrtA-LPETG with peptide outside binding pocket.** AlphaFold3 models are shown for wild-type saSrtA with either a LPETGG substrate sequence (top) or AQALPETGG substrate sequence (bottom). For all, saSrtA is shown in gray cartoon, with the catalytic residue (H120, C184, R197) side-chains as sticks and colored by heteroatom (C=yellow, O=red, N=blue, S=gold). The peptide substrates are in stick representation and colored by heteroatom (C=blue, O=red, N=blue). The substrate is not properly positioned in the active site when only LPETGG was used as the sequence, evidenced by a relatively far distance between C184 and the P1 Thr/P1’ Gly positions, the P4 Leu not binding in its hydrophobic pocket (as in the bottom models), and relatively high variability in overall peptide position. The output model 0 is shown for each to the right.

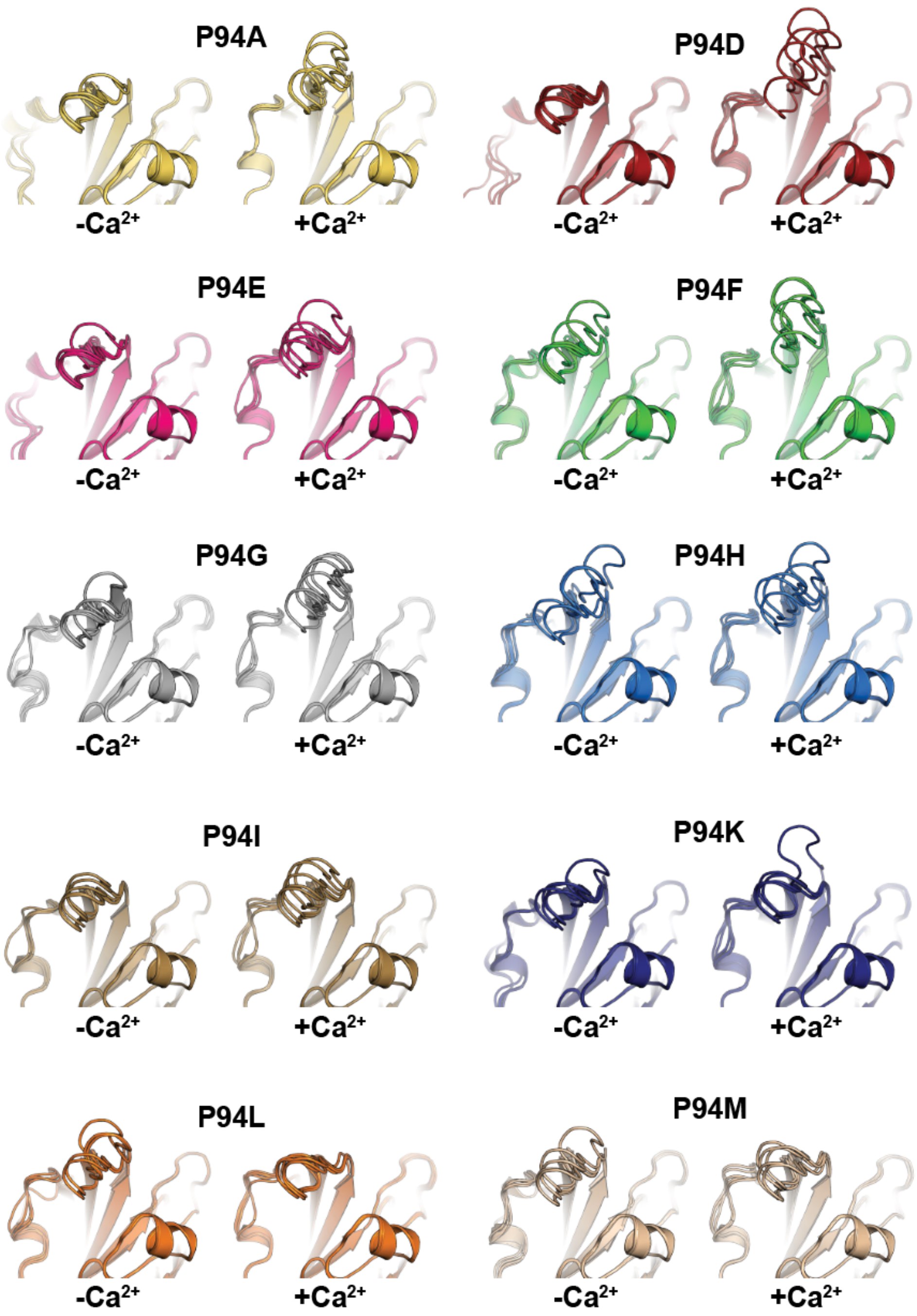
P94A
-Ca2+
+Ca2+
P94D
-Ca2+
+Ca2+
P94E
-Ca2+
+Ca2+
P94F
-Ca2+
+Ca2+
P94G
-Ca2+
+Ca2+
P94H
-Ca2+
+Ca2+
P94I
-Ca2+
+Ca2+
P94K
-Ca2+
+Ca2+
P94L
-Ca2+
+Ca2+
P94M
-Ca2+
+Ca2+

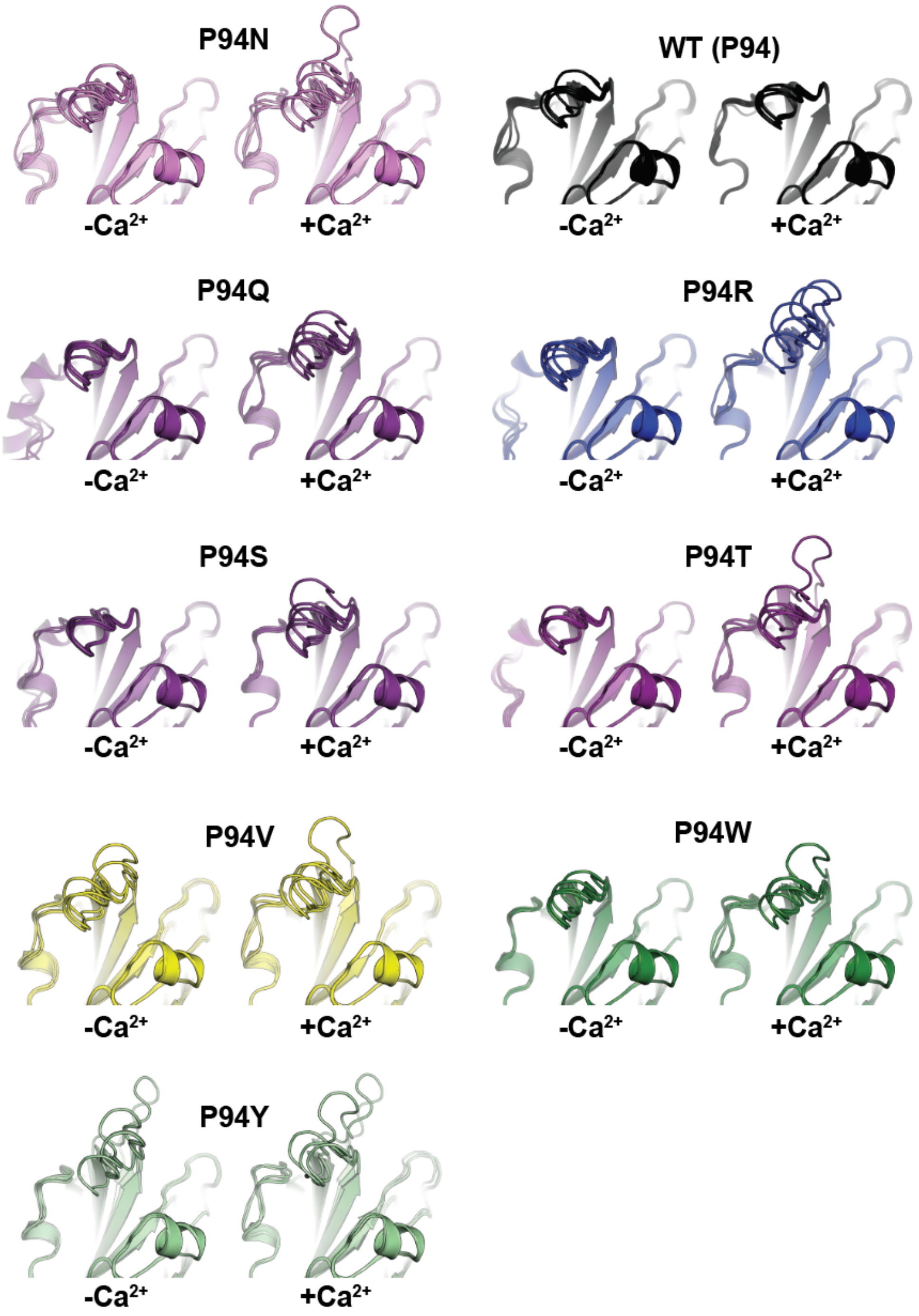

**Figure S5. AlphaFold3 models of P94X saSrtA variants, ± $Ca^{2+}$.** All models are shown in cartoon representation and colored by P94X saSrtA variant. All 5 output models for each variant/condition (± $Ca^{2+}$) were aligned and the relative variability in β7-β8 loop is shown.

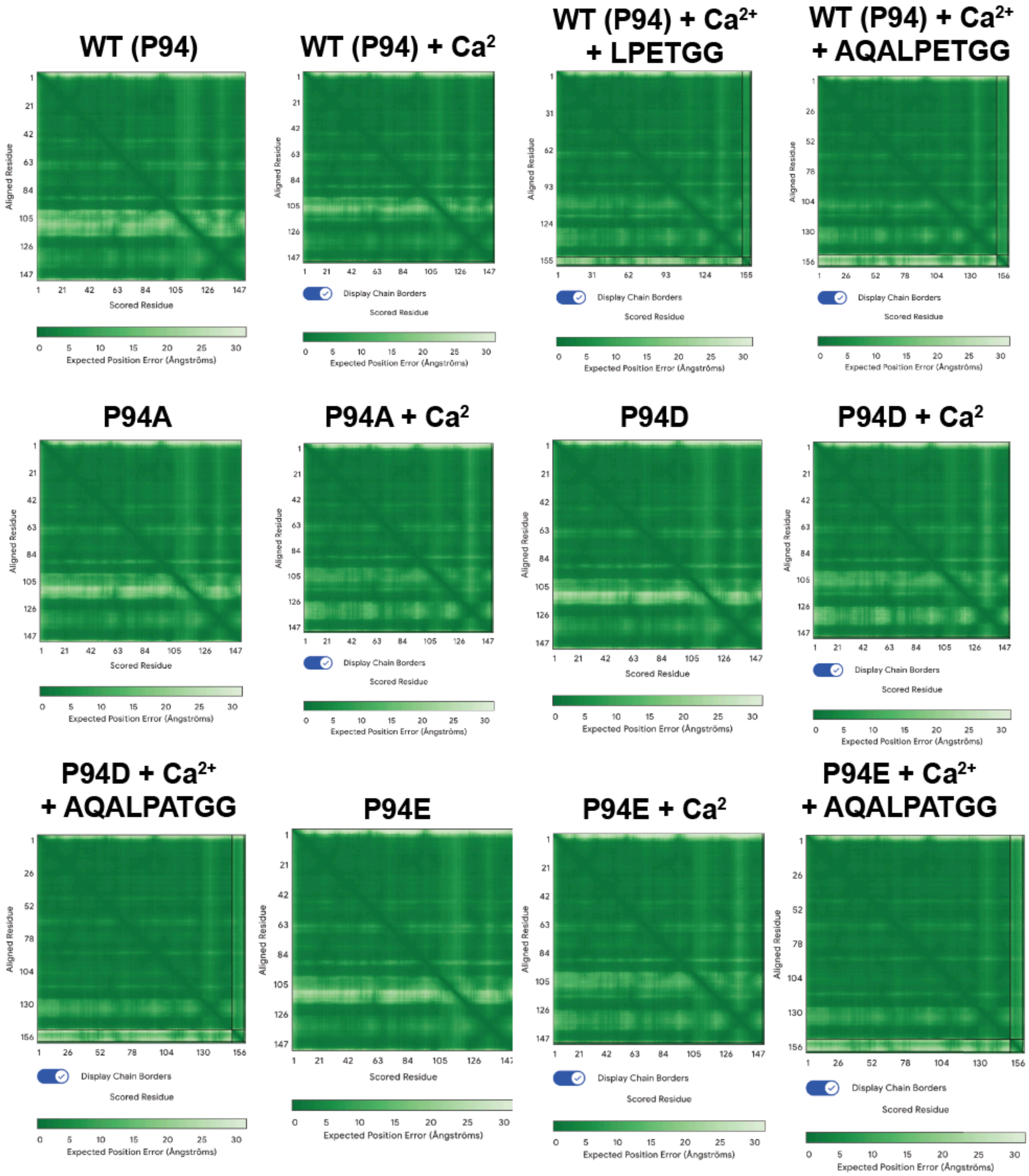
WT (P94)
WT (P94) + Ca2
WT (P94) + Ca2+ + LPETGG
WT (P94) + Ca2+ + AQALPETGG
P94A
P94A + Ca2
P94D
P94D + Ca2
P94D + Ca2+ + AQALPATGG
P94E
P94E + Ca2
P94E + Ca2+ + AQALPATGG
Aligned Residue
Scored Residue
Display Chain Borders
Expected Position Error (Ångströms)

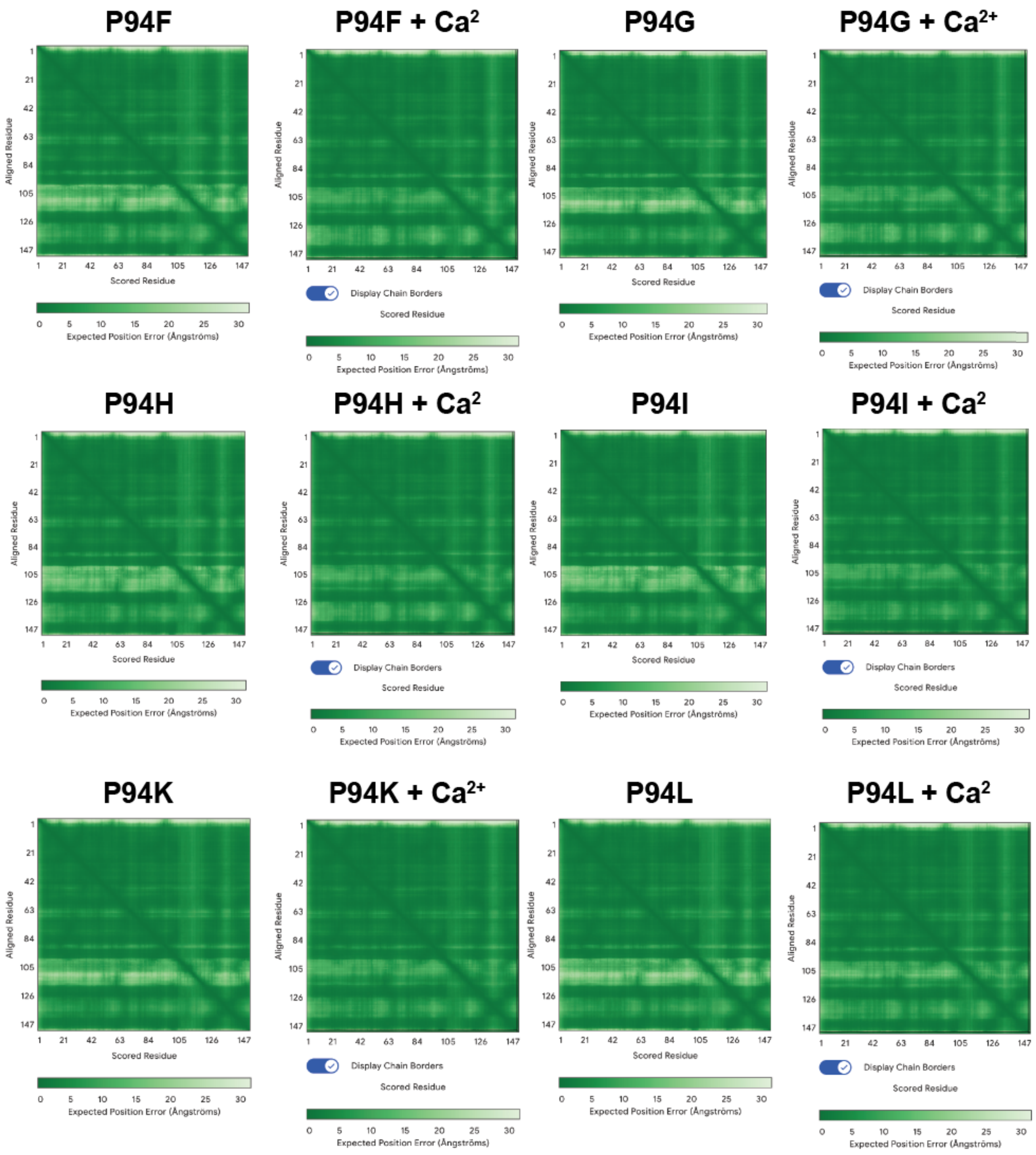
P94F
P94F + Ca2
P94G
P94G + Ca2+
P94H
P94H + Ca2
P94I
P94I + Ca2
P94K
P94K + Ca2+
P94L
P94L + Ca2
Aligned Residue
Scored Residue
Display Chain Borders
Expected Position Error (Ångströms)

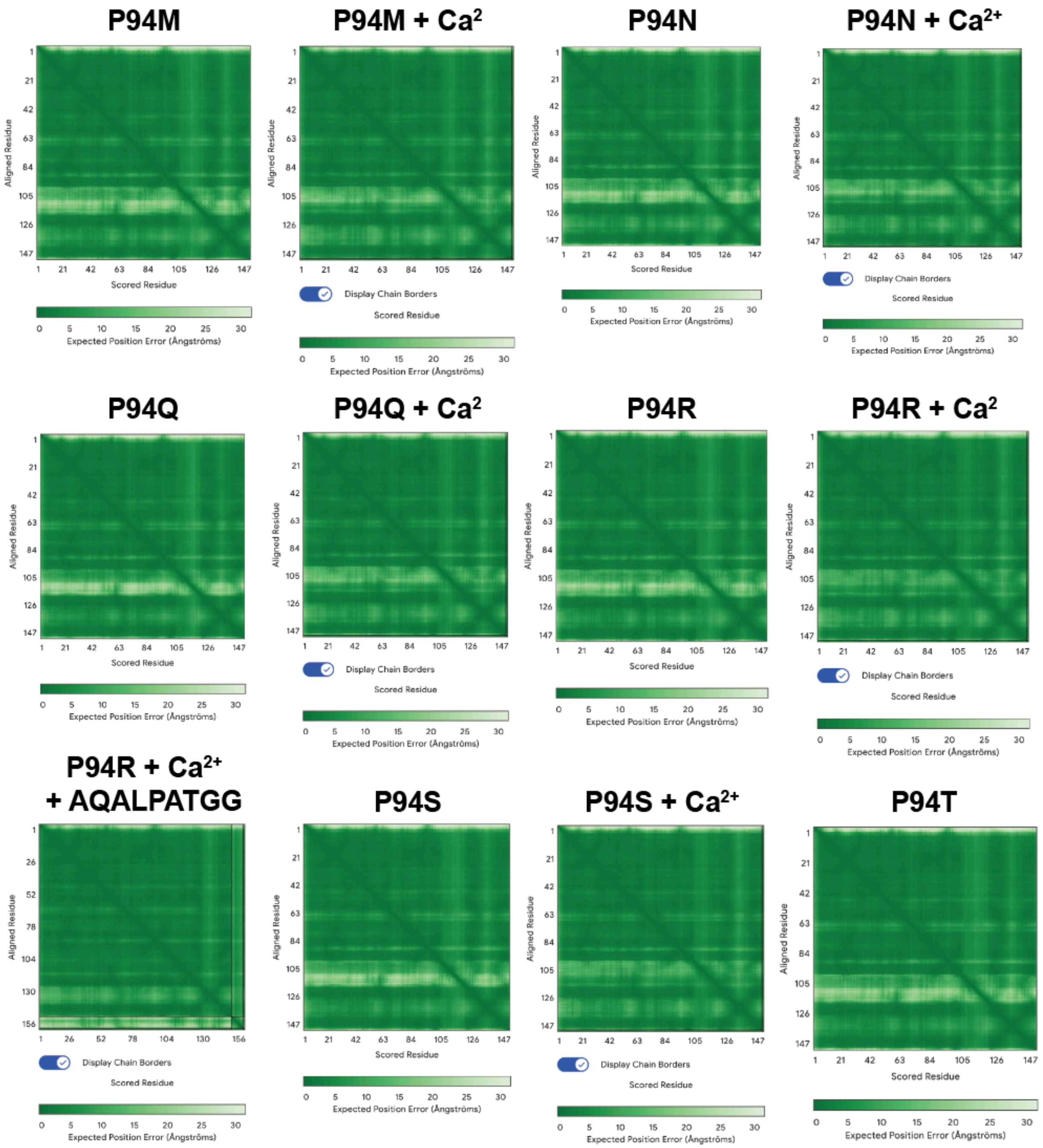
P94M
P94M + Ca2
P94N
P94N + Ca2+
P94Q
P94Q + Ca2
P94R
P94R + Ca2
P94R + Ca2+
+ AQALPATGG
P94S
P94S + Ca2+
P94T
Aligned Residue
Scored Residue
Display Chain Borders
Expected Position Error (Ångströms)
0
5
10
15
20
25
30

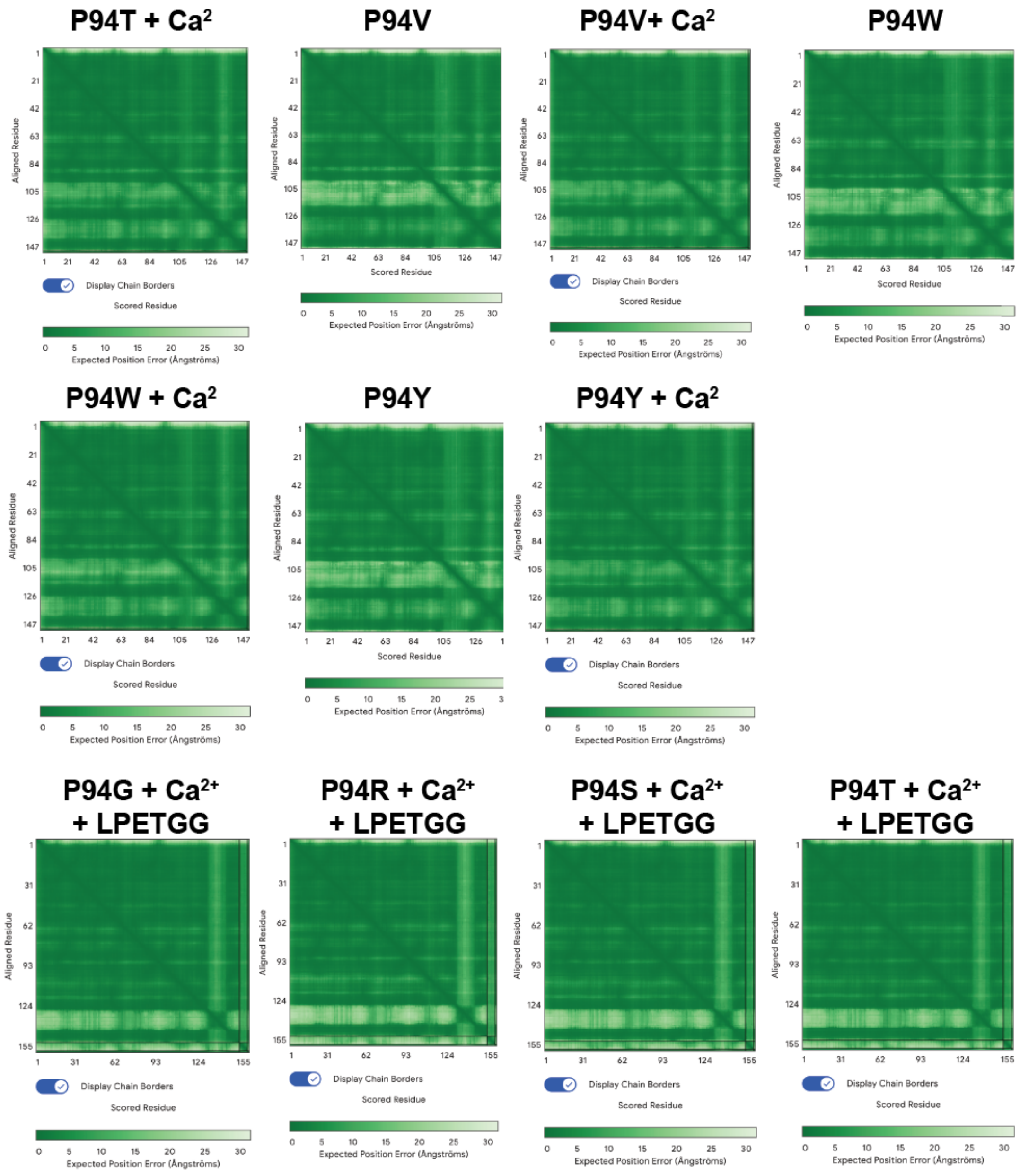

**Figure S6. AlphaFold3 output data.** Output AlphaFold3 data to highlight the quality of the models analyzed. The corresponding output values (pTM, ipTM) are in **Table S2**. The P94G, P94S, and P94T + $Ca^{2+}$ + LPETGG data are shown in **Figure S3A**. These models were very similar to the wild-type (WT) + $Ca^{2+}$ + AQALPETGG models, and alignment of the saSrtA enzymes for output model 0 as compared to WT was RMSD = 0.1 Å (P94G), 0.1 Å (P94S), and 0.1 Å (P94T).

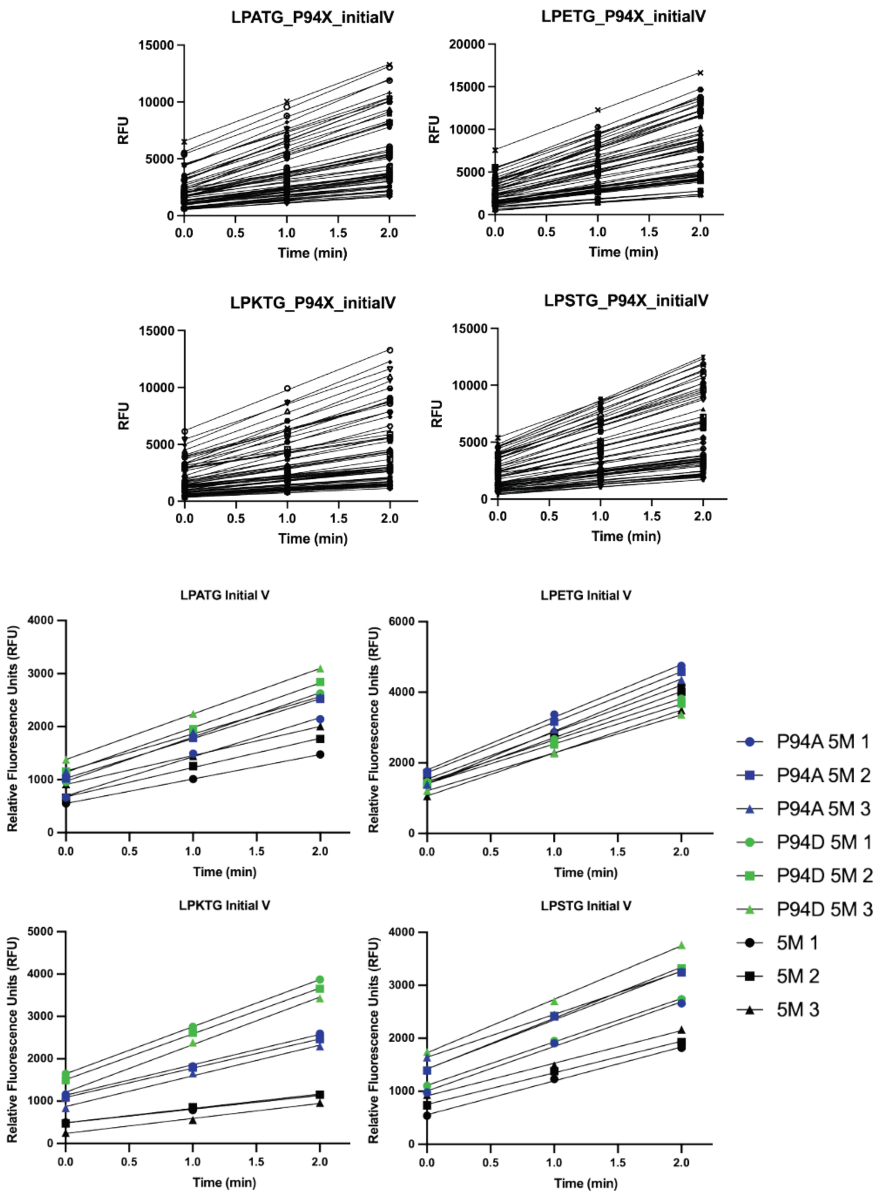

**Figure S7. Initial velocity calculations for P94X saSrtA variants.** The first 3 time points from fluorescence cleavage assays were used to calculate initial velocities for all enzyme-substrate pairs reported. Averaged values ± standard deviations are reported in **Table S4**.

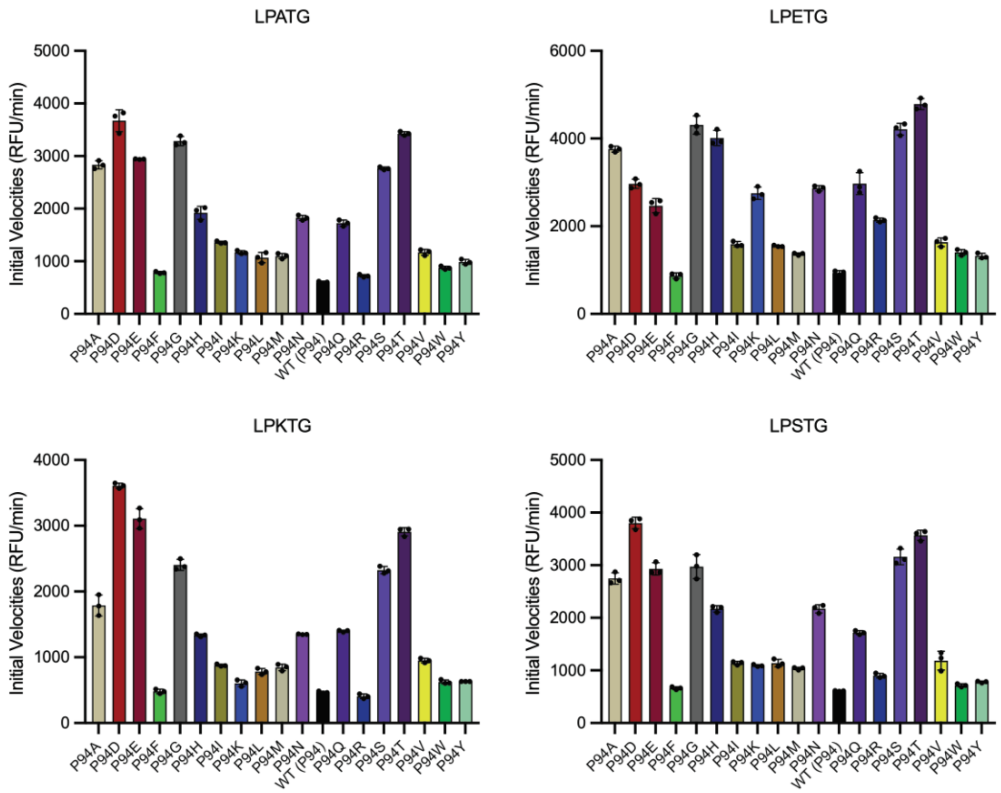

**Figure S8. Comparison of initial velocities for P94X saSrtA variants**. Initial velocities (RFU/min) are shown as the averaged values from triplicate assays ± standard deviation for four peptide substrates: LP**A**TG, LP**E**TG, LP**K**TG, and LP**S**TG. The trends observed are very similar to those for the RFU values at t=20 min (**Figures 2B, 4**).

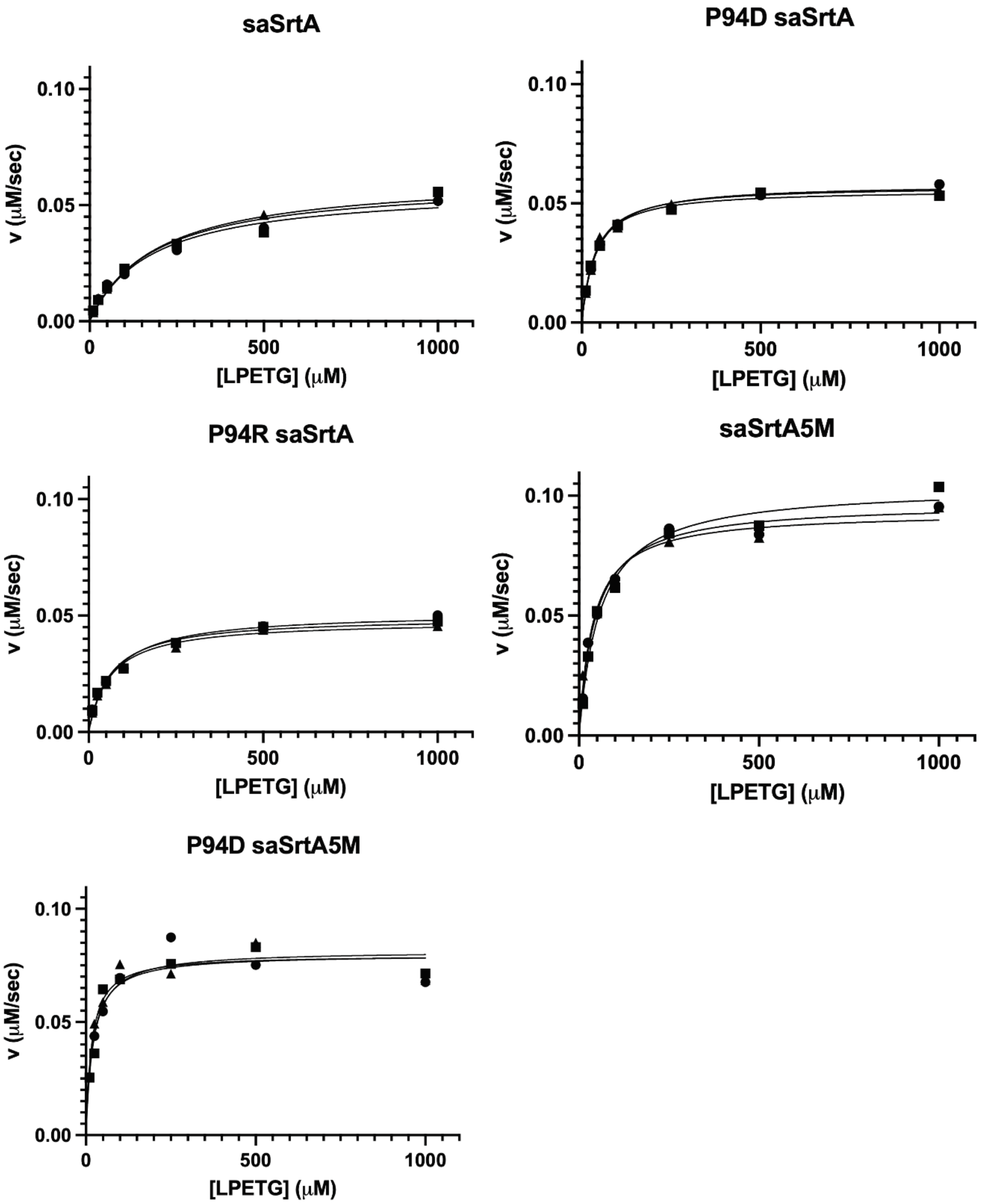

**Figure S9. Replicate data for enzyme kinetics assays.** Replicate data for enzyme kinetics assays (averaged data in **Figure 11A**). Reaction rates were calculated by determining the percentage of product formed at t=10, 60, 120, and 180 s using an HPLC assay. Rates were then determined using a linear regression curve in Excel. All $R^2$ values were >0.98. Parameters ($V_{max}$ and $K_m$) were determined using the Michaelis-Menten fit in GraphPad Prism. For $k_{cat}$, this value was determined using the equation $V_{max} = k_{cat}$*[Enzyme]$_{total.}$

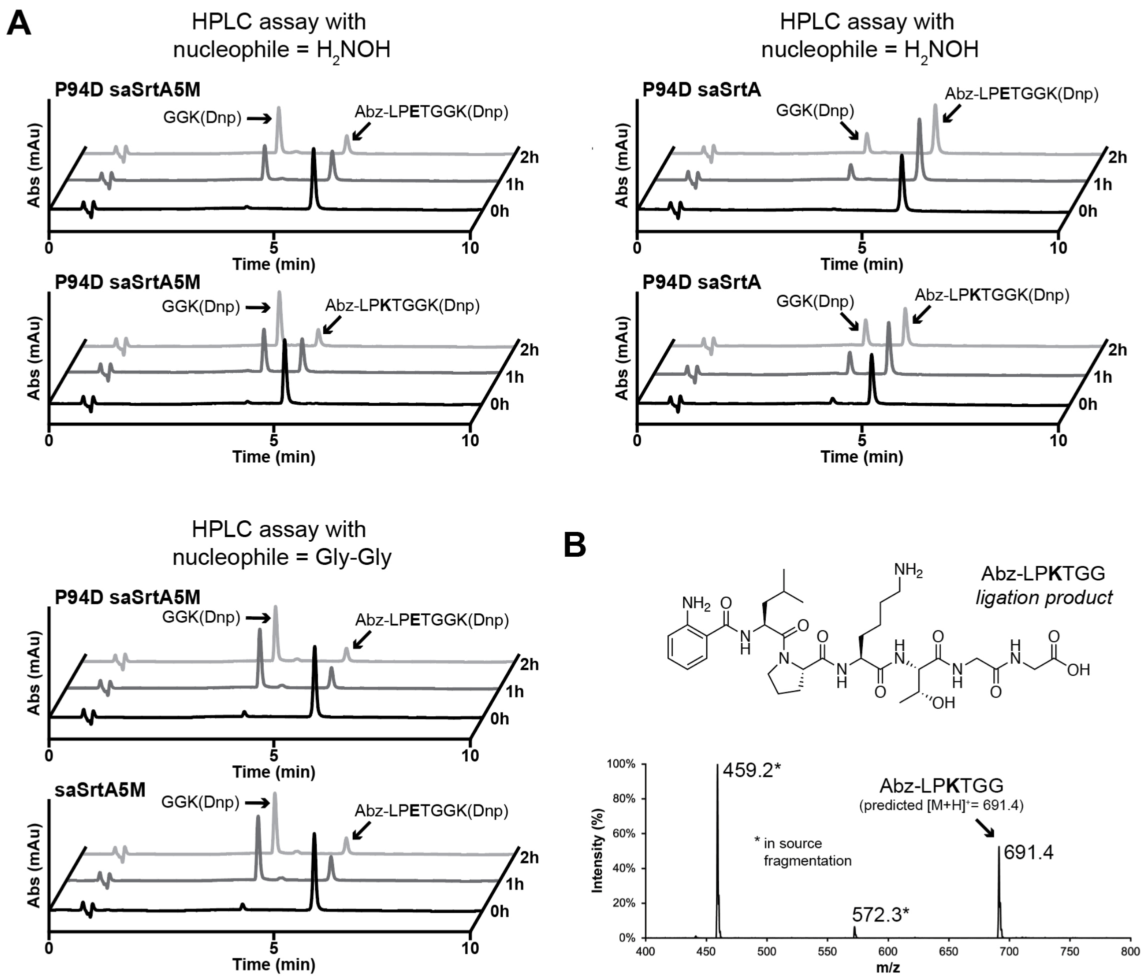

**Figure S10. Representative HPLC traces (360 nm) for assays with saSrtA5M, P94D saSrtA5M, and P94D saSrtA ($H_2NOH$ and/or Gly-Gly nucleophiles)**. (**A**) Representative HPLC traces of sortase-mediated ligations ($H_2NOH$ or Gly-Gly nucleophiles) that show product formation over time, which can be calculated by integration of the indicated peaks (Abz-LP**X**TGGK(Dnp) and GGK(Dnp)). (**B**) Representative mass spectrum for the ligation product (Abz-LP**K**TGG) formed from the reaction of Abz-LP**K**TGGK(Dnp) and Gly-Gly in the presence of P94D SrtA5M, including the molecular structure of the product.
